# Engineered Channel Asymmetry Extends Hydrogen-Bonding Networks for Proton Conduction

**DOI:** 10.64898/2026.04.11.717926

**Authors:** Nolan P. Jacob, Vincent T. Silverman, Gisselle Prida Ajo, Huong T. Kratochvil

## Abstract

The precise and selective transport of protons across cellular membranes relies on the dynamic formation and dissipation of hydrogen-bonding networks involving water molecules, protein sidechains, and backbone carbonyls. As in aqueous solution, protons are conducted over long distances along chains of hydrogen-bonded water molecules within narrow protein pores. To engineer proton-conductive pathways, therefore, we must explicitly account for the dynamic behavior of these networks. In previous work, we showed that incorporation of polar Gln residues into hydrophobic pores drives formation of transient, single-file water wires that enable proton-selective transport. Here, we sought to enhance conduction by introducing targeted Ile-to-Ser substitutions to extend connectivity across the pore. We find that the position of Ser relative to Gln modulates sidechain dynamics and, in turn, channel hydration. Although increased polarity reduces hydrophobic length and enhances hydration, these effects alone do not explain the observed conduction rates. Instead, asymmetry in the arrangement and dynamics of polar sidechains emerges as a key determinant of proton conductivity. Together, these results demonstrate that proton conduction is governed not only by pore polarity and hydration, but also by the dynamic and asymmetric organization of hydrogen-bonding networks. This work establishes design principles for engineering proton-selective channels and reveals how asymmetry enables efficient proton transport across biological membranes.

**Significance Statement:** Proton transport, central to many biocatalytic and bioenergetic processes, requires exceptional selectivity, yet the governing principles remain elusive and difficult to disentangle in natural systems. Designed minimalist proton channels offer new avenues to isolate and test features hypothesized to influence proton conduction. Here, we engineered new-to-nature proton channels with increasingly polar vestibules to establish how control of the hydrogen-bonding network tunes proton conductivity. We found that the effective hydrophobic length, pore polarity, and pore hydration do not contribute significantly to proton conduction rates. Instead, we show that asymmetric sidechain dynamics are key to extending the hydrogen-bonding networks necessary for rapid proton translocation. Our results reveal new tunable parameters that must be considered in the design of proton-selective systems.

## Introduction

Proton channels are responsible for a variety of biological functions including pH regulation(1, 2), ATP synthesis(3), cellular signaling(4), and substrate transport(5). At physiological conditions, the concentration of protons is approximately 10^6^-fold lower than that of other cations like Na^+^ and K^+^. This large disparity necessitates highly selective proton transport to discriminate against competing ions.

The selective conduction of protons is achieved via the Grotthuss shuttling mechanism, in which effective long-range proton transport occurs along hydrogen-bonded networks of water molecules and ionizable functional groups, enabling rapid translocation without net movement of heavy atoms(6–10). Specifically, these dynamic hydrogen-bonded water networks, which often form transient water wires that act as conduits for proton transfer, have been observed in abiological systems including carbon nanotube porins(11, 12), metal organic nanotubes(13), and hydrogen-bonded organic frameworks(14), as well as in native proton channels like gramicidin A(15, 16), influenza A matrix protein(17, 18), cytochrome c oxidase(19–21) and voltage-gated Hv1(22, 23). Thus, controlling the formation and organization of these labile hydrogen-bonding networks is essential for defining proton-conductive behavior in these systems. In the absence of ionizable functional groups, this requires precise arrangement of water molecules into conductive hydrogen-bonding networks(24, 25). However, the fundamental principles that govern how these networks form and support proton conduction remain elusive. Critically, there is no predictive framework linking how specific molecular features control the organization, stability, and connectivity of these hydrogen-bonding networks.

*De novo* membrane protein design has enabled the creation of new-to-nature ion channels and transporters(26–31) and is a powerful platform for dissecting fundamental principles of channel structure–function relationships. In the context of proton-selective channels, this approach has provided direct mechanistic insight into the features that govern proton transport. Previous work by Lear *et al*. demonstrated that the low complexity peptide LS2, composed of Leu and Ser in a repeating arrangement of (LSLLLSL)_3_, assembles into tetrameric bundles(32). The polar vestibule created by exclusion of Ser from the lipid environment facilitated the organization of proton-conducting water networks within the pore, supporting proton translocation(33, 34). Recent work by Kratochvil *et al*. established that a dynamic, hydrogen-bonded water wire is the minimal functional unit for proton conduction. In a simple hydrophobic pentameric assembly(35), incorporation of polar layers within the pore created hydrogen-bonding sites that promote pore hydration, stabilize transient water wires, and support proton transport(36). Thus, these simplified *de novo* designed systems offer unique opportunities to directly interrogate the molecular determinants of proton conduction by isolating and tuning specific interactions within a controlled structural framework.

Here, we leverage functional *de novo* proton channels to directly define how pore hydration and sidechain dynamics govern hydrogen-bonding networks and, ultimately, proton conductivity. We increase pore polarity by systematically introducing Ser residues at key pore-lining positions (**Fig. 1**), with the rationale that shortening the hydrophobic length lowers the energetic barrier for forming hydrogen-bonded water networks required for proton conduction. We find that, although pore polarity, and consequently pore hydration, are tunable, they are not sufficient on their own to control proton conductivity. We demonstrate that sidechain dynamics and channel asymmetry are designable features that must be explicitly considered in the design of functional proton-selective channels.

**Figure 1.**
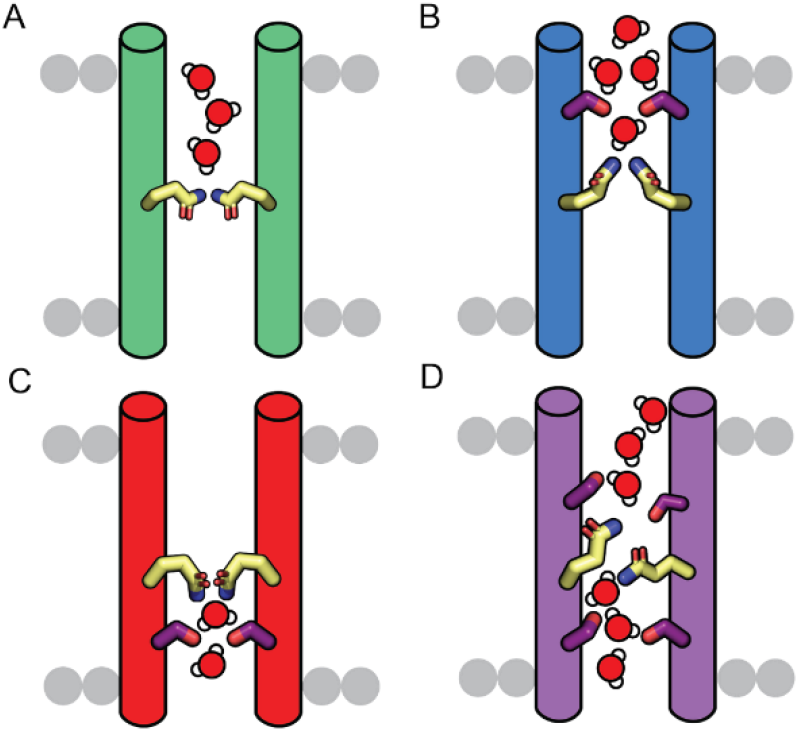
Extending the hydrogen-bonding network in proton-conductive channels. (A) Parent variant with single polar Gln in pore. (B) Designed channel with Ser substitution one helical turn above the Gln. (C) Designed channel with Ser substitution one helical turn below the Gln. (D) Designed channel with two Ile-to-Ser substitutions.

## Results

### Ile-to-Ser substitutions increase pore polarity while maintaining pentamer assembly

In our original design scaffolds, LQLL and LLQL, Gln occupies the *a* positions of the heptad repeat at positions 10 and 17, respectively (**Fig. 2A,B**). The hydrophobic Ile at the *d* positions, located one helical turn above and below the Gln residues, are ideal sites for introducing polar substitutions that can extend the hydrogen-bonding network within the pore. To increase pore polarity, we selected the smallest polar residue, Ser. In principle, Ile-to-Ser substitutions should add polar functionality to our channels with minimal perturbation of pentamer assembly.

**Figure 2.**
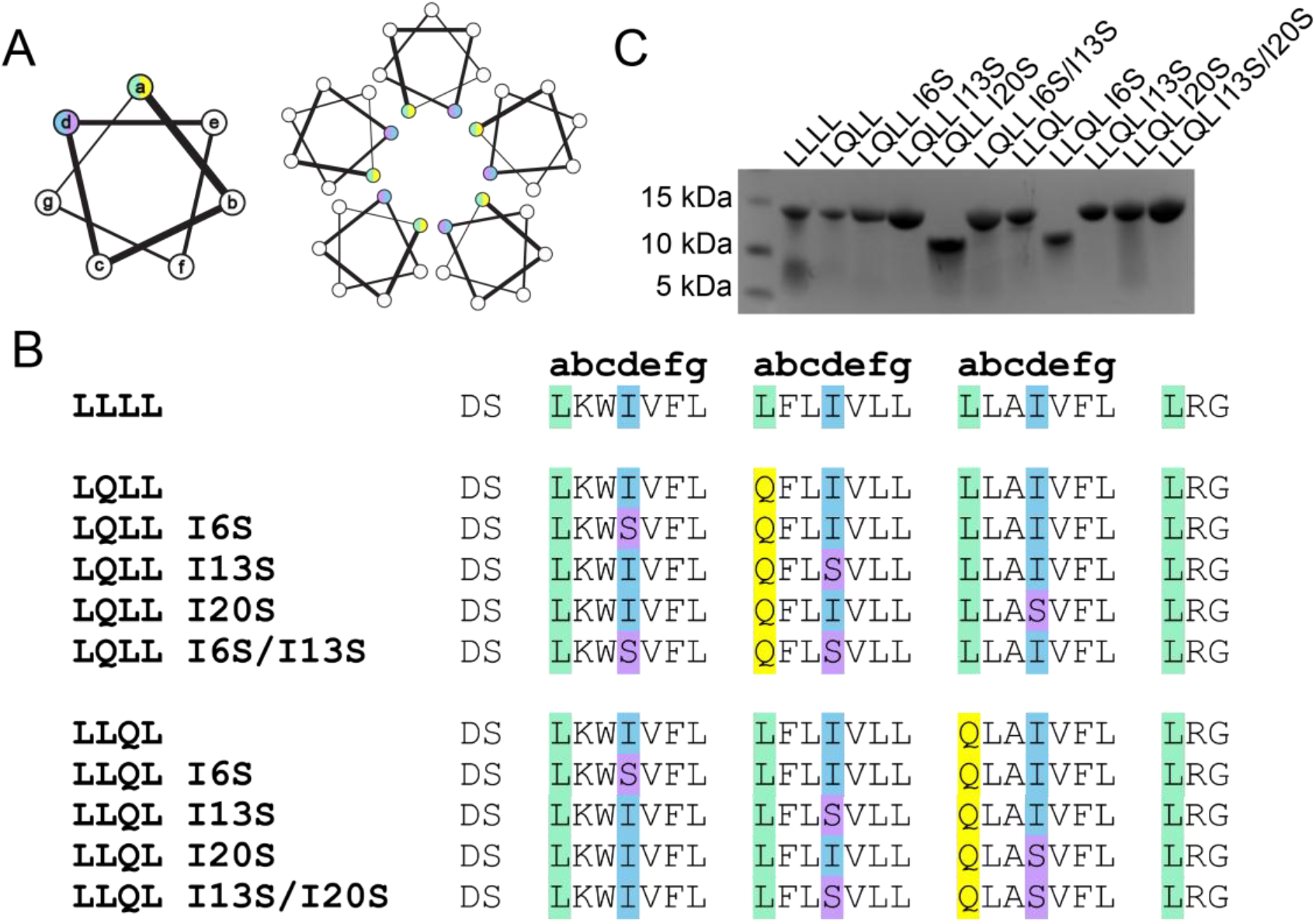
Ile-to-Ser mutant sequences. (A) Cartoon representation of helical wheel with *a* and *d* pore-facing positions colored. (B) Sequences for LLLL, original designs LQLL and LLQL, and all Ser mutants. Pore-facing residues are highlighted accordingly: Leu (green), Ile (blue), Gln (yellow), and Ser (purple). (C) SDS-PAGE of all peptides. All peptides except LQLL I120S and LLQL I6S formed pentameric assemblies.

In this study, we mutated the Ile residues at the *d* positions to Ser, producing two series of Ile-to-Ser peptides (I6S, I13S, and I20S) from the parent scaffolds (**Fig. 2B**). We also generated select double mutants with Ile-to-Ser substitutions at both *d* positions flanking the Gln layer in the pore (LQLL I6S/I13S and LLQL I13S/I20S).

All designed peptides were synthesized using solid-phase peptide synthesis and purified using established workflows (see **Materials and Methods, SI Appendix Table 1**). To assess whether the Ile-to-Ser substitutions affected pentamer assembly, we analyzed the peptides by SDS-PAGE, which we previously showed can serve as a rapid proxy for determining membrane peptide oligomerization state(35, 36). LLLL, LQLL, and LLQL form defined pentamers with bands at ∼ 15kDa (**Fig. 2C**, lanes 1, 2, and 7). LQLL mutants with a Ser positioned one helical turn above or below the Gln (*e*.*g*., LQLL I6S or LQLL I13S) also form pentamers (**Fig. 2C**, lanes 3 and 4). The double Ser mutant, LQLL I6S/I13S, which combines both substitutions, similarly assembles into a pentamer (**Fig. 2C**, lane 6). In contrast, the I20S mutant, where the Ser lies more than two helical turns away from the Gln, migrates as a tetramer (**Fig. 2C**, lane 5). For the LLQL series, we observe the same trend: the Ile-to-Ser substitutions near the Gln residue (*e*.*g*., I13S, I20S, and the double mutant I13S/I20S) do not disrupt pentamer assembly (**Fig. 2C**, lanes 9-11) and Ile-to-Ser substitutions distal to the Gln (LLQL I6S) abolish pentamer formation (**Fig. 2C**, lane 8).

Gel electrophoresis showed that Ile-to-Ser substitutions minimally perturb pentamer assembly when introduced at *d* positions one helical turn from the Gln residue. These LQLL (I6S, I13S, I6S/I13S) and LLQL (I13S, I20S, I13S/I20S) mutants are consequently more polar than their parent scaffolds, allowing us to directly test how increased channel polarity affects proton conduction.

### Double Ile-to-Ser substitutions show increased proton conduction rates relative to the parent scaffold

To determine the effects of the Ser substitutions on proton conduction rates, we measured channel activity using established liposomal assays (**Fig. 3A**)(36–38). Asymmetric liposomes were prepared with excess internal K^+^ and excess external Na^+^ (see **Materials and Methods**). Addition of the K^+^ carrier valinomycin allows K^+^ to flow down its concentration gradient out of the liposomes, creating an electrochemical potential that drives protons into the liposomes through channels or carriers. Protonation of the encapsulated dye, 8-hydroxypyrene-1,3,6-trisulfonate (HPTS), decreases its fluorescence, which we measure and convert to total proton concentration using calibration curves (**SI Appendix Fig. S1**). Beyond distinguishing proton-selective from non-proton-selective channels, this assay also enables direct comparison of proton conduction rates among our variants.

**Figure 3.**
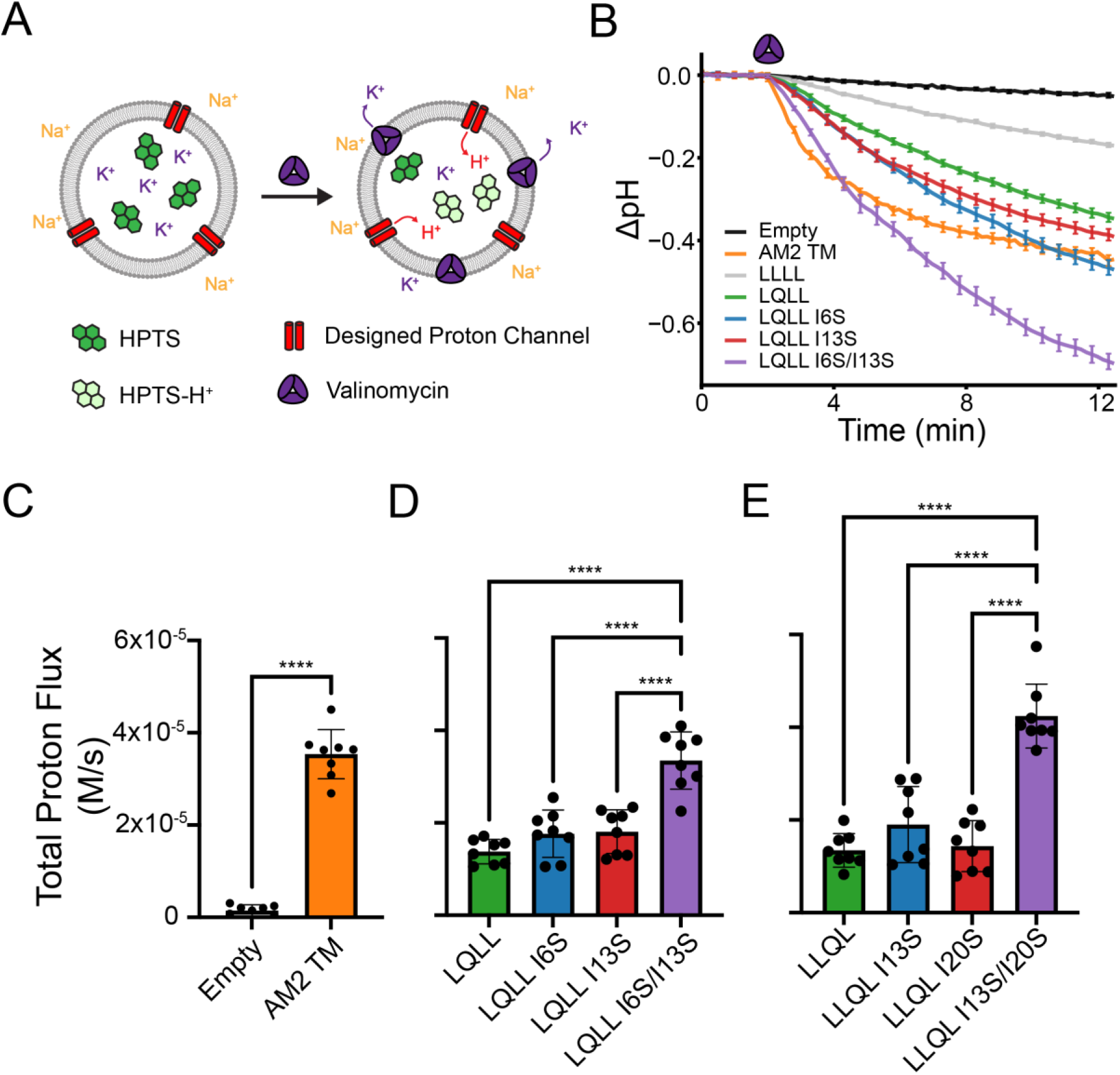
Proton flux liposomal assay. (A) Cartoon schematic of the liposomal assay. (B) Raw data for controls and LQLL family plotted in terms of ΔpH. Error bars represent the standard error of the mean across all 8 biological replicates. (C-E) Initial rates of total proton flux for controls and the LQLL and LLQL mutants. (C) The positive control, AM2-TM, showed significant proton selective transport activity against the negative empty control (unpaired *t*-test, p<0.00001). (D) Single Ser mutants show no significant difference from the parent LQLL in proton conduction rates, whereas the double mutant LQLL I6S/I13S differs significantly from all other variants. The data was analyzed using one-way ANOVA and post-hoc Tukey’s test. (E) Similarly, single Ser mutants show no significant difference from the parent LLQL in proton conduction rates, whereas the double mutant LLQL I13S/I20S differs significantly from all other variants. The data was analyzed using one-way ANOVA and post-hoc Tukey’s test.

The time-dependent change in ΔpH for the controls and design variants is shown in **Fig. 3B**. Valinomycin (**Fig. 3A,B**, purple) is added at t = 2 min, driving proton flux into the liposome and resulting in a decrease in intraliposomal pH. As expected, empty liposomes, which contain no protein, show no change in ΔpH upon addition of the K^+^ carrier. In contrast, liposomes containing the natural proton channel from the Influenza A virus, AM2 TM, show a significant decrease in ΔpH (**Fig. 3B**). To compare initial rates of proton flux, we fit the slope of the ΔpH curves over the first 60 seconds following addition of valinomycin and corrected the rates to account for buffering (see **Materials and Methods**). Comparing initial total proton flux rates in **Fig. 3C**, we can see a clear difference between our negative control (empty) and our positive control (AM2 TM) (**SI Appendix Table 2**).

Initial rates for LLLL, a hydrophobic non-proton-conducting channel, and the LQLL and LLQL parent designs are consistent with prior experimental data (**SI Appendix Fig. S2, Table 3**). Comparing proton conduction rates for LQLL and its Ser variants, we find that the single Ser mutants (LQLL I6S and LQLL I13S) exhibit rates comparable to the parent LQLL (**Fig. 3D, SI Appendix Table 4**). One-way ANOVA and post-hoc Tukey’s tests indicate no significant differences in proton conduction rates between LQLL and the single Ser mutants (see **Materials and Methods**). In contrast, the double mutant LQLL I6S/I13S shows significantly higher proton conduction rates than the parent and both single Ser mutants.

A similar pattern emerges with LLQL designs (**Fig. 3E, SI Appendix Table 5**). Initial rates of the single Ser mutants LLQL I13S and LLQL I120S show no significant difference compared to the LLQL parent. The double mutant LLQL I13S/I20S exhibits significantly higher initial proton flux rates than the parent LLQL and the single Ser mutants (**Fig. 3E**).

### Increasing pore polarity alone does not enhance proton conduction

Our liposomal assays reveal that the Ser variants retain proton-selective activity. Interestingly, single Ser substitutions within the conduction pathway do not affect proton conduction rates, despite increasing the polarity of the channel. Notably, the double Ser mutants show enhanced proton transport activity compared to both parent and single Ser mutants. These results suggest that factors beyond simple channel polarity contribute to determining proton conduction rates.

To probe the effects of the Ile-to-Ser substitutions with atomic-level detail, we performed classical molecular dynamics (MD) simulations for pentamer-forming variants identified by SDS–PAGE. Three independent 200-ns simulations were carried out for each variant in 1-palmitoyl-2-oleoyl-sn-glycero-3-phosphocholine (POPC) bilayers.

When we consider the effective hydrophobic lengths of our variants (**SI Appendix, Fig. S3, Table 6**), no systematic relationship emerges with proton conduction. Notably, introducing the Ile-to-Ser substitution at position 6 in LQLL increased the effective hydrophobic length of the channel, defined as the longest hydrophobic span calculated from MD simulations of each variant. When we incorporated Ser one helical turn below the Gln residue, as in LQLL I13S, the effective hydrophobic length decreased, but the addition of a second Ser in the double mutant LQLL I6S/I13S did not further reduce this length (**SI Appendix Fig. S3**). Although the effective hydrophobic lengths of these variants are similar, their proton conductivities differ significantly (**Fig. 3D**). A similar trend is seen for LLQL I13S and LLQL I13S/I20S, which both have effective hydrophobic lengths of ∼12 Å, yet the double mutant exhibits significantly higher proton conduction rates (**Fig. 3E** and **SI Appendix Fig. S3**). These data suggest that simply minimizing the effective hydrophobic length does not alter proton conductivity, implying that additional factors are involved.

### Incorporation of Ser alters pore hydrogen-bonding networks

The introduction of polar Ser reshaped the hydrogen-bonded network, notably expanding hydrogen-bonding interactions throughout the pore. To characterize these changes, we used Bridge2(39) to examine the dynamic hydrogen-bonding networks from representative simulations (**SI Appendix Fig. S4**).

Each node is a hydrogen-bonding residue (*e*.*g*., Gln or Ser), and each edge represents a hydrogen-bonding interaction between two residues, with up to five water molecules mediating the interaction. Edges have been filtered to require an occupancy greater than 10% of the 1000 frames in each simulation. Indeed, the additional polar Ser increased the hydrogen-bonding interactions between the different layers. In the parent LQLL and LLQL (**SI Appendix Fig. S4**), the hydrogen-bonding interactions are largely restricted to the Gln layer, with both direct and water-mediated interactions linking Gln sidechains. The addition of a single Ser above or below the Gln (**SI Appendix Fig. S4**) extended the network across the polar sidechains over an average of 2.8 or 2.3 water molecules, respectively. With both Ser in the pore, this network spans three layers and an average of 3 water molecules. The enhanced polarity from the incorporation of Ser does increase the hydrogen-bonding network in the pore, but proton channel activity does not scale linearly with the number of polar residues.

The types of hydrogen-bonding interactions in the pore change drastically with the introduction of Ser (**SI Appendix Fig. S5, Table 7**). For the parent LQLL and LLQL, Gln–Gln hydrogen bonds account for >50% of all hydrogen-bonding interactions within the pore. With the Ser one helical turn above the Gln, as in LQLL I6S and LLQL I13S, we observe a shift in hydrogen-bonding partners, with an increase in Gln–Ser interactions and a decrease in Gln–Gln interactions (**SI Appendix Fig. S5**). In contrast, LQLL I13S and LLQL I20S, with the Ser one helical turn below, show an increase in Gln–Gln hydrogen bonding, which now accounts for >80% of all interactions in the pore (**SI Appendix Fig. S5**). Finally, in the double mutants LQLL I6S/I13S and LLQL I13S/I20S, Gln–Gln interactions are minimized, with a corresponding increase in hydrogen bonding between Gln and nearby water molecules (**SI Appendix Fig. S5**). Indeed, the redistribution of hydrogen-bonding interactions may contribute to the observed changes in proton flux. We note that the double mutants, which exhibit significantly higher proton conduction rates than both the parent and single Ser variants, show the lowest fraction of Gln–Gln hydrogen-bonding interactions (<20%) and the highest fraction of Gln–water interactions (>30%) (**SI Appendix Fig. S5**). Our findings suggest that the types of hydrogen-bonding interactions within the pore play a key role in modulating proton conduction. Increased interactions between Gln sidechains and water molecules appear to promote the formation of more continuous and dynamic hydrogen-bonding networks, which are likely critical for efficient proton transport.

### Ile-to-Ser substitutions alter Gln sidechain conformational dynamics and channel hydration

We further analyzed our simulations to assess how Ile-to-Ser substitutions influence pore dynamics and hydration. Although classical MD simulations do not explicitly model proton transfer, they enable characterization of the structural organization and dynamics of hydrogen-bonding networks that govern proton transport. To examine the Gln sidechain dynamics, we defined 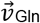, a vector extending from the Cα to the Nε of the Gln sidechain (**Fig. 4A**). For each trajectory, we extracted the coordinates of 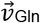 for the five Gln residues in the pentamer at every snapshot (**Fig. 4B**) and plotted their orientations. Additionally, we extracted the χ_1_ and χ_2_ Gln sidechain dihedral angles for all Gln in every simulation frame to identify the dominant rotameric states sampled in each trajectory (see **Materials and Methods**). We also evaluated how the increased polarity from the Ile-to-Ser substitutions affected channel hydration using the Channel Annotation Package (CHAP)(40).

**Figure 4.**
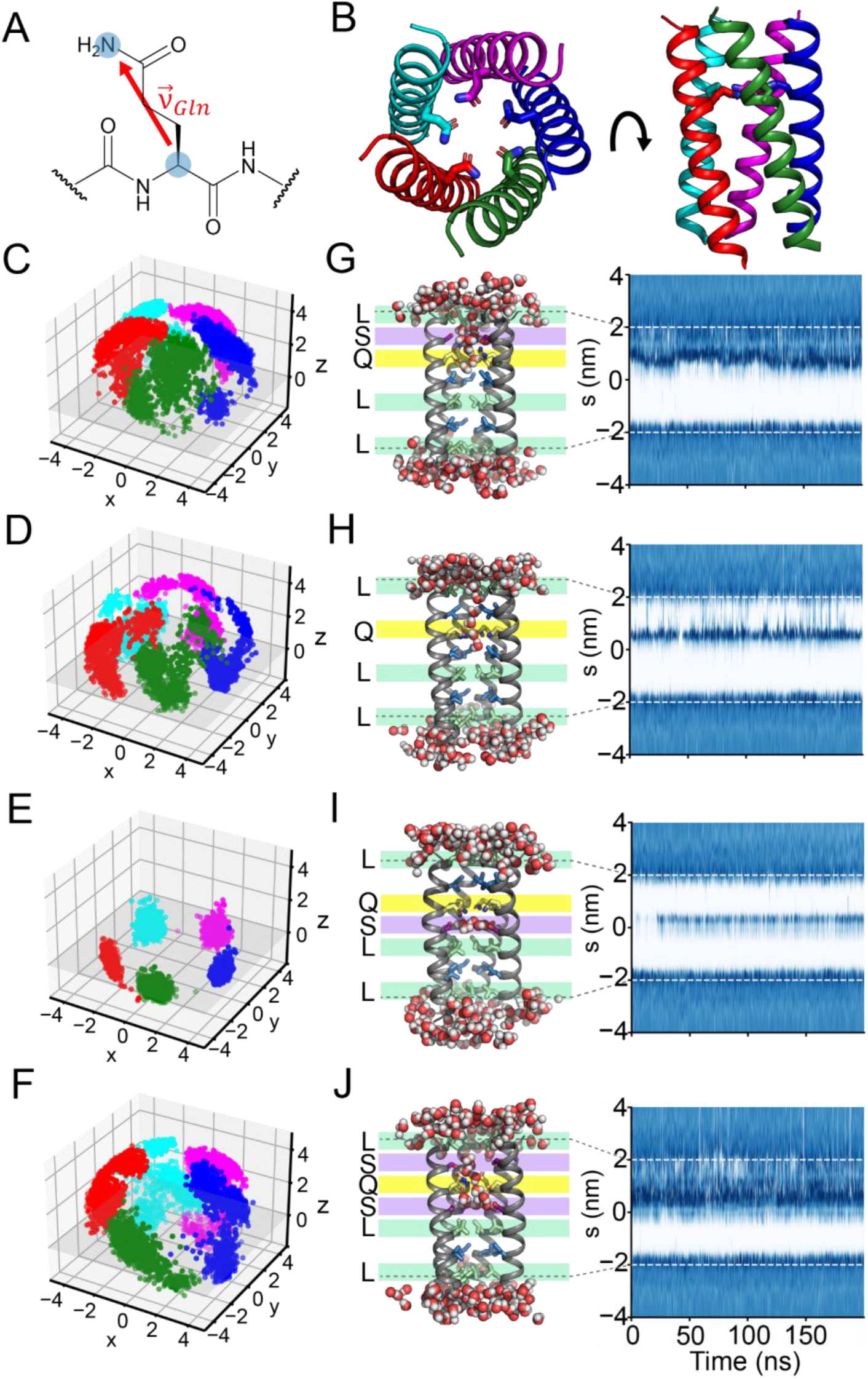
MD simulations indicate Ser substitutions affect Gln sidechain dynamics and pore hydration. (A) 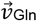 is defined as the vector from the Cα to the Nε of the Gln sidechain. (B) Structure of LQLL (pdb 7UDZ) with each of its five helices colored to denote the separate chains. For each of the mutants (C) LQLL, (D) LQLL I6S, (E) LQLL I13S, and (F) LQLL I6S/I13S, 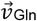 is plotted for the five Gln10 residues over the course of the simulation. The Cα plane is denoted in gray for (x, y, z = 0). Representative snapshots and water density profiles from a single production run for (G) LQLL, (H) LQLL I6S, (I) LQLL I13S, and (J) LQLL I6S/I13S are shown. Waters within 3 Å of the channel are shown, and the pore-facing residues of interest are colored as in **Fig. 1**. Dashed lines represent the channel boundaries within the simulation.

For LQLL variants (**Fig. 4C-F**), we observe notable changes in the Gln sidechain conformations when we incorporate Ser residues at the *d* positions. For the parent LQLL, we found that the Gln sidechain samples many conformational states, with most 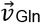 oriented above the plane defined by the five Gln Cα atoms (**Fig. 4C, SI Appendix Fig. S6**). Analysis of χ_1_ and χ_2_ sidechain dihedrals from the simulations shows two major clusters that account for 99% of sampled conformations (**SI Appendix Fig. S7**). These clusters correspond to orientations centered at an average position of z = 1.24 Å above the Cα plane (**SI Appendix Table S8**).

When we introduced the Ile-to-Ser substitution one helical turn above in LQLL I6S, we noted a shift in the orientations of 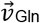 (**Fig. 4D, SI Appendix Fig. S8**). Compared to the parent LQLL, the 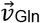 distribution for LQLL I6S is shifted toward more positive z-values, with an average position of z = 3.09 Å above the Cα plane (**SI Appendix Table S8**), indicating that the Gln sidechain more frequently points upward along the pore axis. Looking at the χ_1_ and χ_2_ dihedral angles, we discover a major cluster that constitutes 81.1% of the conformations and a minor cluster that constitutes 17.2% (**SI Appendix Fig. S9**), both of which have positive z-values for 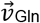.

When the Ser is one helical turn below the Gln, as in LQLL I13S, the Gln sidechain conformations are markedly altered, becoming restricted below the Cα plane (**Fig. 4E, SI Appendix Fig. S10**). Here, the 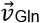 distribution is highly localized around z = -0.39 Å (**SI Appendix Table S8**). Notably, we see one major cluster that accounts for 99% of all Gln conformations in our χ_1_ and χ_2_ dihedral angle analysis (**SI Appendix Fig. S11**). In almost all frames, we see that the Gln is locked into a conformation where 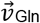 has a negative z-value.

When both flanking *d* positions were mutated to Ser in LQLL I6S/I13S, 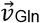 distributions again span a wide range of *z*-values (**Fig. 4F, SI Appendix Fig. S12**). Here, we see three conformations in our χ_1_ and χ_2_ dihedral analysis. The major conformational cluster, which captures 53.5% of the Gln conformations, is defined by rotameric states with z = 3.21 Å (**SI Appendix Fig. S13, Table S8**). This cluster corresponds to the major cluster identified in the LQLL I6S simulations. The second largest cluster accounts for 31.0% of the Gln conformations and includes states corresponding to the major conformation observed in LQLL I13S, where the 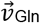 is below the Cα plane. 14.1% of the Gln conformations sampled in MD simulations of LQLL I6S/I13S fall into a third cluster defined by χ_1_ ≈ -150º and χ_2_ ≈ 50º dihedral angles, resulting in sidechain rotamers with positive z-values (**SI Appendix Fig. S13**).

We uncovered a similar effect on the Gln sidechain conformations for the LLQL series (**SI Appendix Fig. S14**). For the parent LLQL, we see that 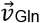 vectors lie near the Gln Cα plane (**SI Appendix Fig. S15**). Consistent with this, analysis of the χ_1_ and χ_2_ dihedral angles reveals a single dominant conformational cluster comprising 95.8% of sampled states (**SI Appendix Fig. S16, Table S8**), in which the 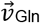 z-values are centered around 1.19 Å. When we make a single Ser substitution to the Ile13 position, we see the 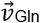 vectors shift upward along the z-axis (**SI Appendix Fig. S17**), clustering around 3.3 Å above the Cα plane. Again, we observe two dominant clusters that together comprise 99.2% of all Gln conformations (**SI Appendix Fig. S18, Table S8**). For LLQL I20S, where the Ser is one helical turn below the Gln residue, we again see constrained Gln sidechain dynamics with 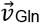 distributions centered around z = -0.4 Å (**SI Appendix Fig. S19**). As in the LQLL I13S mutant, we observe a single major conformation (95.8%), characterized by 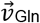 vectors with negative z-values (z = -0.2 Å) (**SI Appendix Fig. S20, Table S8**). For the double mutant LLQL I13S/I20S, 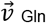 vectors indicate that the Gln sidechains sample a broad range of conformations (**SI Appendix, Fig. S21**). Here, we see three major clusters of Gln conformational states. The first cluster, which contains 71.7% of the Gln rotamers sampled, is identical to the major clusters from LLQL I13S and LQLL I6S (**SI Appendix, Fig. S22, Table S8**). The second-most populated cluster, comprising 14.2% of the Gln conformations, is centered around χ_1_ ≈ −75° and χ_2_ ≈ −50°, and includes conformations positioned just above the Cα plane. The third cluster, representing 4.1% of the Gln states, has average z-values of about z = 0.5 Å.

The Ile-to-Ser substitutions also affected the distribution of water within the channel, as reflected in the distinct hydration profiles of each variant (**Fig. 4G-J**). We reproduced the hydration profiles for the parent scaffold LQLL (**Fig. 4G, SI Appendix Fig. S6**). In both cases, we see that the water near the polar Gln site occasionally forms short-lived water wires that connect to bulk water at the top or bottom of the channels, respectively. Incorporating the Ser substitution one helical turn above the Gln in LQLL I6S resulted in water occupancy throughout the upper third of the channel (**Fig. 4H, SI Appendix Fig. S8**). In contrast, adding Ser one helical turn below the Gln in LQLL I13S resulted in decreased water density throughout the pore (**Fig. 4I, SI Appendix Fig. S10**). Snapshots reveal a pocket of stable, long-lived water (**SI Appendix Fig. S23**) in ring-like hydrogen-bonded structures between the Gln10 and Ser13 layers of the pore. Unlike LQLL and LQLL I6S, where channel water molecules flicker and interact with bulk water outside the pore, the hydration profile of LQLL I13S suggests that channel waters in this variant are confined. Finally, the water density profile for the double mutant (LQLL I6S/I13S) shows that more than half of the pore remains hydrated throughout the 200-ns simulation (**Fig. 4J, SI Appendix Fig. S12**). Our simulations of LQLL I6S/I13S reveal dynamic water networks that exchange between the channel and bulk.

MD simulations of the LLQL series corroborate our findings from the LQLL series, with analysis revealing consistent trends in water organization despite slight differences in hydration profiles. For LLQL I13S, we see increased hydration between the Ser13 and Gln17 layers of the channel (**SI Appendix Fig. S14H, S17**). Because the Ser13 layer lies at the pore center, hydration remains localized and does not extend across the full pore. Like LQLL I13S, when the Ser is directly below the Gln in the helix, in LLQL I20S, we see a marked decrease in pore water density (**SI Appendix Fig. S14I, S19**). Again, we observe the persistence of ring-like water structures that do not appear to exchange with the bulk water below the channel. With LLQL I13S/I20S, we discover increased water occupancy in the bottom two-thirds of the channel near the polar residues (**SI Appendix Fig. S14J, S21**).

Our MD simulations reveal that introducing polar Ser residues one helical turn from the Gln modulates the Gln sidechain conformational dynamics, altering both its orientational distribution and the conformational states within the pore. Furthermore, the increased polarity of the channel influences pore hydration. When Ser is positioned above the Gln layers, we observe increased water density within the channel. In contrast, when Ser lies below the Gln (e.g., LQLL I13S or LLQL I20S), water occupancy in the pore decreases.

### Crystal structures reveal distinct Gln conformations and pore symmetry

We solved lipidic cubic phase X-ray crystal structures for several of our designs, one for each positional variant of the Ser: LLQL I13S, with the Ser one helical turn above the Gln (**Fig. 5B**); LQLL I13S, where Ser is below the Gln (**Fig. 5C**); and the double Ser mutant LQLL I6S/I13S (**Fig. 5D**). All three structures formed distinct pentameric assemblies that are less than 0.55 Å RMSD of the parent design. As in LQLL (pdb 7UDZ, **Fig. 5**), electron density in the pore of LLQL I13S and LQLL I6S/I13S can be modeled as water. Looking at the hydrogen-bonding structures within the pore (**Fig. 5A-D**, bottom), we note that the Gln conformations are highly symmetric in all but the LQLL I6S/I13S structures. This symmetry is also apparent when we extract the χ_1_ and χ_2_ dihedral angles of the Gln sidechains in the crystal structures (**SI Appendix Table S9**).

**Figure 5.**
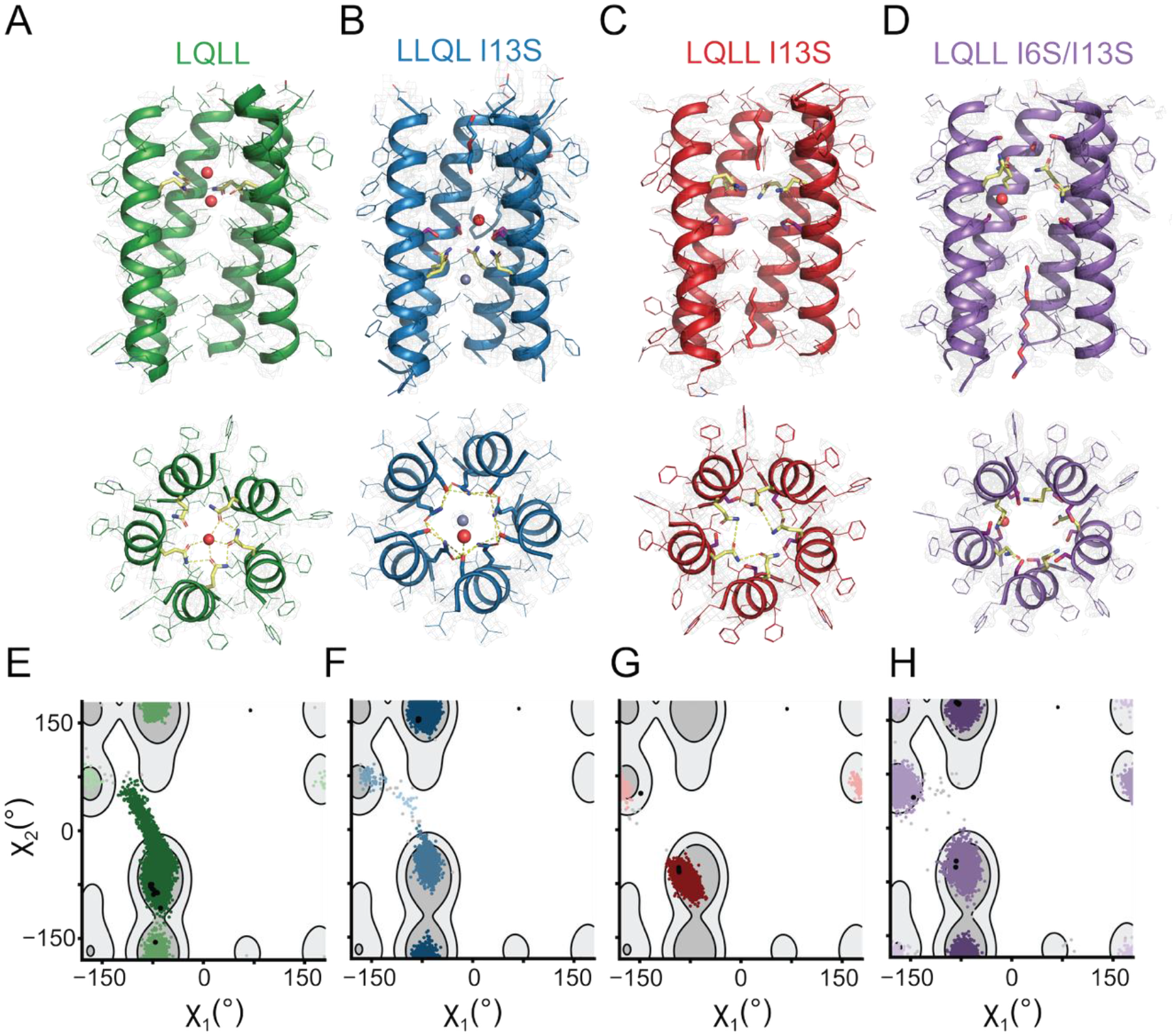
X-ray structures and Gln rotamer analysis of Ser mutants. Side-on views and top-down views of crystal structures of (A) LQLL (from pdb 7UDZ), (B) LLQL I13S, (C) LQLL I13S, and (D) LQLL I6S/I13S. One helix is hidden for clarity. Pore-facing Ser (purple) and Gln (yellow) residues are highlighted. Waters and other small molecules within the pore are shown. Hydrogen-bonding interactions between crystallographic water and/or sidechains are also shown. (E-H) Plots of χ_1_/χ_2_ dihedral angles from Gln sidechains sampled from MD simulations (colored) and from crystal structures (black) for each of the mutants. These are overlaid on a contour map of Gln rotameric states sampled from a curated structural database of transmembrane proteins, where lighter gray contours denote the top 70% and darker gray contours denote the top 90%. Simulation points are colored by cluster with more prevalent clusters represented in darker colors.

The Gln sidechain dihedrals modeled in our crystal structures are in strong agreement with the conformations observed in MD simulations. In the crystal structure for LLQL I13S (**Fig. 5A**), we see that all Gln sidechains are oriented within hydrogen-bonding distance of the Ser above. Consistent with this, the χ_1_ and χ_2_ dihedral angles for all ten Gln residues in the asymmetric unit fall within the same cluster as the most populated conformation observed in MD simulations of LLQL I13S (**Fig. 5E**). For the LQLL parent, we note that nine of the ten Gln sidechains in the asymmetric unit adopt the same conformation (**Fig. 5B**), which corresponds to the major conformation observed in MD simulations (**Fig. 5F**). One Gln sidechain assumes a conformation within the second-most populated cluster.

In the LQLL I13S crystal structure (**Fig. 5C**), where the Ser is one helical turn below the Gln, all Gln sidechains point downward, and their χ_1_ and χ_2_ dihedral angles fall into the same cluster that accounts for 99% of the frames sampled in MD simulations (**Fig. 5G**). These Gln residues are again positioned within hydrogen-bonding distance of the Ser. In the solved structure of the double mutant LQLL I6S/I13S, we model Gln in several distinct conformational states (**Fig. 5D**). When we compare the sidechain dihedrals with those extracted from MD simulations, our crystal structures corroborate the *in silico* models. MD simulations of this mutant reveal three major conformational clusters, each of which is captured in the crystal structure (**Fig. 5H**). Furthermore, all three conformations are observed in both copies of the pentamer within the asymmetric unit, providing the first experimental evidence for channel asymmetry in the double mutant.

### Polar residue placement defines rotamer diversity within the pore

Our MD simulations (**Fig. 4**) and X-ray crystallographic structures (**Fig. 5**) demonstrate that strategic placement of Ser within the pore modulates Gln sidechain dynamics and defines the set of accessible rotameric states for the Gln residue. By extracting the z-coordinate from 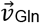 for all five Gln residues across 1000 frames per simulation (15,000 data points per variant), we clearly observe a shift in Gln orientation relative to the Cα plane upon introduction of the Ile-to-Ser substitutions (**Fig. 6A, SI Appendix Fig. S24**).

**Figure 6.**
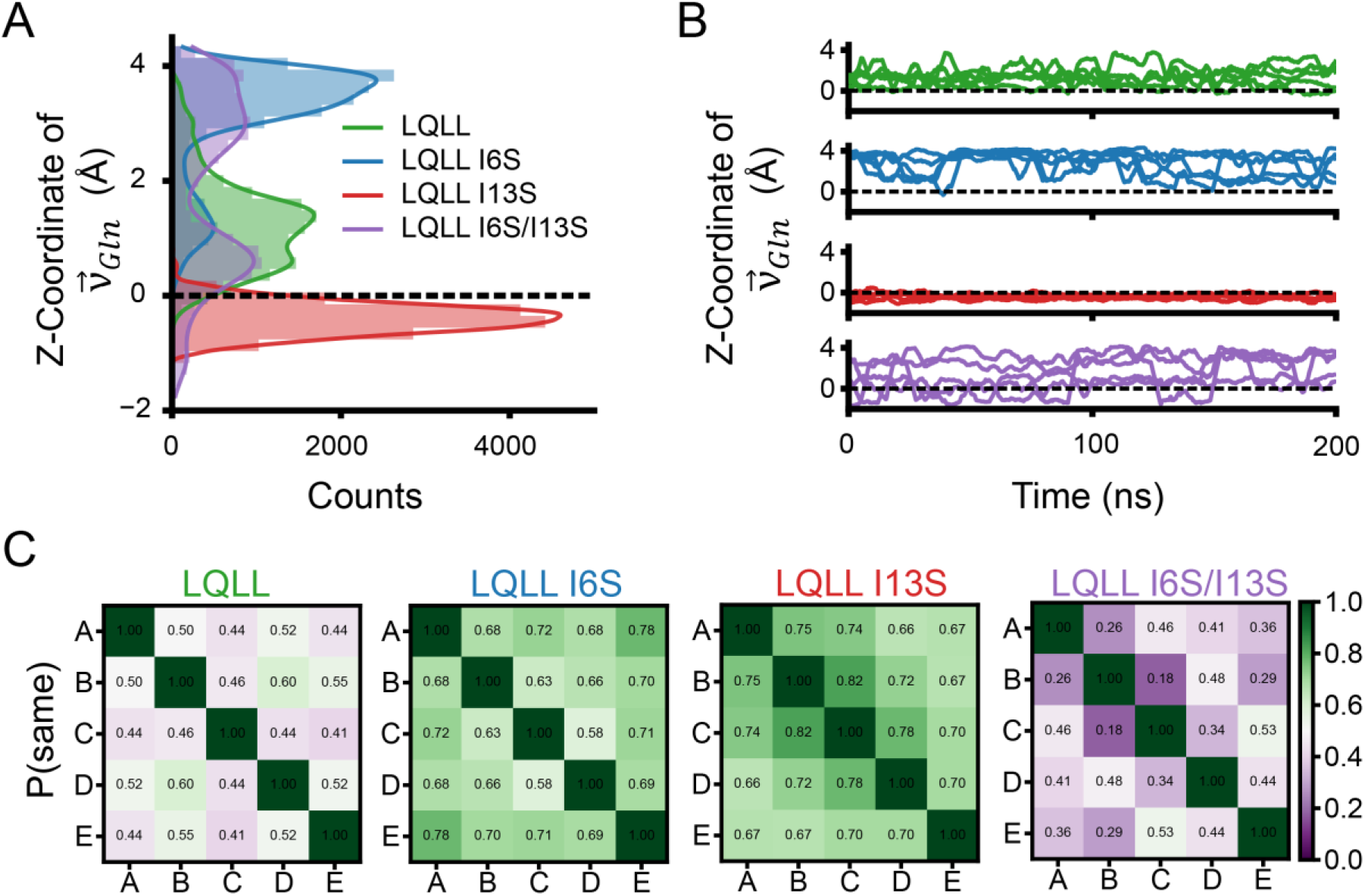
MD simulations reveal double Ser mutant breaks channel symmetry. (A) Histogram of z-coordinate of 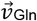 for all Ser mutants of LQLL. Counts include all five Gln10 residues within the pore across the full simulation and all three replicates for a total of 15,000 data points per variant. The Cα plane is shown as a dashed line. (B) Line plots of the z-coordinate of each of the five pore-lining Gln10 residues as a function of time for a single simulation trajectory of each mutant. (C) Pairwise state agreement matrix for Gln sidechain in LQLL variants.

For LQLL, the average z-value for 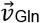 is centered around z = 1.4 Å above the Cα plane (**Fig. 6A**, green). With the single Ser above the Gln, as in LQLL I6S, the Gln sidechain orientation shifted up the channel, with average z-values for 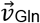 at z = 3.3 Å, accounting for the population distribution (**Fig. 6A**, blue). When the Ser is one helical turn below the Gln, the 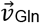 distributions shift dramatically to z = - 0.3 Å, below the Cα plane (**Fig. 6A**, red). With two Ser in the pore, the 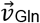 distributions sample a broad range of rotameric states that span from z = -2 to 4 Å along the pore (**Fig. 6A**, purple). Similarly, for LLQL and its Ser variants, we see the same effects: the 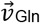 distributions can be tuned by the position of the Ser residue relative to the Gln (**SI Appendix, Fig. S24**).

We demonstrate that Gln sidechain conformations can be tuned by the positioning of Ser within the pore. While these effects on Gln conformations are evident from our analysis, they do not fully explain the differences observed in proton flux rates (**Fig. 3**). We initially hypothesized that distinct rotameric states correspond to functional and nonfunctional conformations, with more constrained states that restrict Gln mobility, like those seen in LQLL I13S or LLQL I20S, expected to be less conducive to proton conduction. However, our results indicate that all sampled conformations remain functional, suggesting that proton conduction is not limited to a single preferred rotameric state.

### Breaking symmetry in Gln orientations extend hydrogen-bond networks

This prompted us to assess how collective Gln sidechain organization, rather than individual rotamer states, governs the formation and connectivity of hydrogen-bonding networks within the pore. To capture collective Gln sidechain dynamics, we extracted the z-coordinate of 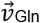 for each of the five pore-lining Gln in every frame for each independent simulation and plotted these values as a function of time (**Fig. 6B**). In the parent LQLL and single Ser mutants (LQLL I6S and LQLL I13S), the Gln sidechains move in a highly coordinated manner. In LQLL (**Fig. 6B**, green), all five residues maintain nearly identical z-values over time. Similarly, in LQLL I6S (**Fig. 6B**, blue) and LQLL I13S (**Fig. 6B**, red), the Gln sidechains adopt the same z-position at each time point, reflecting strongly symmetric motion along the pore axis. In contrast, LQLL I6S/I13S (**Fig. 6B**, purple) breaks this symmetry, with individual frames showing each of the five Gln sidechains occupying different z-positions simultaneously. These observations raise the possibility that pore symmetry influences proton conductivity.

We next examined how symmetric versus asymmetric Gln dynamics could modulate proton conductivity. To quantify symmetry in Gln sidechain dynamics, we constructed pairwise state agreement matrices for all designs (**Fig. 6C**), which report the extent to which Gln residues occupy the same conformational state over time and provide a measure of correlated behavior between Gln sidechains. We classified the z-coordinate of each Gln into three states: down (z < 0.0), neutral (0.0 < z < 2.5), and up (z > 2.5) based on the histogram peaks (**Fig. 6A, SI Appendix Fig. S24**). Pairwise state agreement between Gln residues (chains A–E) was then computed for each frame and averaged over the trajectory. Agreement values near 1 indicate that two Gln residues occupy the same state, whereas values near 0 indicate that they rarely share the same state. The parent LQLL displays moderate agreement (∼0.5), while LQLL I6S and LQLL I13S show higher agreement (>0.5), consistent with more symmetric dynamics (**Fig. 6C**). In contrast, LQLL I6S/I13S exhibits the lowest agreement (<0.5), reflecting increased asymmetry.

Similarly, in the LLQL series, we observe symmetric Gln conformational dynamics for the LLQL parent and the single Ser mutants (**SI Appendix Fig. S24B**). Pairwise state agreement matrices for the LLQL variants reveal that the parent LLQL exhibits modest agreement (∼0.4), consistent with slightly correlated Gln behavior and a degree of asymmetry. In contrast, the single Ser mutants display more strongly correlated Gln sidechain dynamics (>0.5), indicative of more symmetric, concerted behavior (**SI Appendix Fig. S24C**)

For the LLQL I13S/I20S trajectory presented in **Fig. S24C** (purple), we note symmetric, coordinated Gln dynamics in the first ∼100 ns of the simulation and asymmetric dynamics in the latter half of the simulation. Analysis of the LLQL I13S/I20S hydration profiles (**SI Appendix Fig. S26A**) suggests that pore hydration may play a role in inducing asymmetry in the pore. To test this hypothesis, we pooled all simulation frames based on pore hydration. Dry frames were classified as any frame with n = 0 or 1 waters as these frames had no wires of waters forming within the pore. Wet frames were defined as any frames after an n ≥ 10 threshold was met to allow for slight fluctuations in the number of waters. (**SI Appendix Fig. S26B**). When the pore is dehydrated, we note that the Gln are symmetric in their dynamics (**SI Appendix Fig. S26C**). This is also evident in the pairwise state agreement matrix, which shows values exceeding 0.9 (**SI Appendix Fig. S26D**). The wet pore tells a different story, with the Gln sidechains adopting asymmetric conformations (**SI Appendix Fig. S26C**). Analysis of the pairwise state agreement matrix reveals that the sidechain dynamics become very asymmetric in the presence of water (**SI Appendix Fig. S26E**). Furthermore, analysis of hydrogen-bonding interactions within the pore reveals that, in the wet state, Gln–Gln interactions are minimized while Gln–water hydrogen bonds are maximized, accounting for >50% of all hydrogen bonds (**SI Appendix Fig. S26F**). This further supports our finding that the type of hydrogen-bonding interactions, rather than their overall abundance, plays a critical role in governing proton conduction. Here, LLQL I13S/I20S provides a unique test case for linking pore hydration to Gln dynamics. In the absence of water, the Gln sidechains exhibit highly symmetric dynamics, and hydrogen-bonding interactions are dominated by Gln–Gln and Gln–Ser contacts. Upon hydration, Gln–water interactions disrupt this symmetry, leading to more asymmetric Gln dynamics. These results highlight pore hydration and asymmetric channel dynamics as critical for extending hydrogen-bonding networks that enable efficient proton conduction.

## Discussion

In this study, we introduced Ile-to-Ser substitutions into the conduction pathway of minimalist proton channels to enhance proton conduction rates. We hypothesized that increasing pore polarity through incorporation of small, polar Ser residues would be sufficient to promote formation of the dynamic hydrogen-bonded water networks required for proton translocation. Our combined computational and experimental results reveal that, while pore polarity influences channel hydration, it is insufficient to modulate proton conductivity. To extend these networks within the pore, we must look beyond water dynamics and explicitly design protein sidechain dynamics and channel asymmetry.

Indeed, prior work in both biological and abiological proton-conducting systems has shown that sidechain conformations and their accessibility can modulate proton transport and conduction mechanisms(41–44). For example, Roy *et al*. demonstrated that sidechain flexibility in bioinspired materials enables these sidechains to adopt configurations that facilitate proton transfer between adjacent layers(44). More broadly, the dynamics of both protonatable and non-protonatable sidechains within hydrogen-bonding networks have been shown to modulate proton flux and selectivity(45).

Asymmetric conformations within otherwise symmetric ion channels can control ion selectivity and conduction(46–48). Similarly, proton conduction pathways are often dynamically regulated, such that local asymmetries in sidechain behavior and hydration determine when and where conductive networks form. For example, work on respiratory complex I demonstrates that proton transfer pathways arise at symmetry-related locations but are gated by protonation-dependent hydration and conformational changes, resulting in functional asymmetry within an otherwise symmetric architecture(49). Asymmetry is also a designable parameter, as demonstrated by Zhou et al., who showed that asymmetric sidechain dynamics of Arg residues govern gating and anion selectivity in designed ion channels(31). In the design of proton- and ion-conductive materials, asymmetric distributions of functional groups and broken symmetry in electrochemical potential have been shown to give rise to rectification, gating, and selective ion transport(50, 51). Here, we demonstrate that the same guiding principles also govern proton conduction in biological pores.

Taken together, these findings establish sidechain dynamics and asymmetry as key design principles for controlling proton conduction. They reveal that proton conductivity depends not only on pore hydration and polarity but also on the dynamic organization of the conduction pathway, and introduce new, tunable parameters for the rational design of synthetic channels. More broadly, these results underscore the importance of these features across both biological and abiological proton-conductive systems.

## Materials and Methods

### Classical MD Simulations

The coordinates of LLLL (PDB ID: 6MCT) were used as the starting structure for all protein models. CHARMM-GUI(52–54) was used to assemble the full simulation systems. During the PDB manipulation step, point mutations (*e*.*g*., L10Q, I13S) were introduced at the appropriate positions, and the C-terminus was modified to an amide to match experiments. The helical bundles were then oriented in the membrane using the Orientations of Proteins in Membranes tool for Positioning Proteins in Membranes (PPM 2.0)(55). A lipid bilayer composed of 1-palmitoyl-2-oleoyl-glycero-3-phosphocholine (POPC) was constructed with an initial 1:1 distribution across leaflets, yielding ∼72–75 lipids per leaflet. Water thickness was set to 2.5 nm. The system was neutralized and brought to 0.15 M KCl. All components were parameterized with the CHARMM36m force field(56), and GROMACS input files were generated. Simulations were performed at 298.15 K.

Minimization, equilibration, and production runs were performed using GROMACS 2021.5(57). All simulations used the recommended CHARMM36 cut-offs. Electrostatic interactions were treated with the Particle Mesh Ewald settings with a Coulomb cutoff of 1.2 nm, and Van der Waals interactions were force-switched between 1.0 and 1.2 nm. Steepest descent energy minimization was performed for up to 5000 steps, until the maximum force was less than 1000 kJ mol^−1^ nm^−1^. Following minimization, a 125 ps NVT equilibration was carried out with position restraints applied to backbone atoms, sidechains, and lipids. Temperature was maintained at 298.15 K using the velocity-rescale thermostat with a coupling constant of 0.1 ps. Subsequently, a 15 ns NPT equilibration was performed with Cα atoms restrained, using the semi-isotropic Berendsen barostat (pressure of 1 bar, compressibility of 4.5 × 10^-5^ bar^-1^, coupling constant of 5 ps) and the Berendsen thermostat (temperature of 298.15 K, coupling constant at 0.001 ps).

Unrestrained production MD simulations were performed for 200 ns, with trajectories saved every 20 ps. Temperature was maintained at 298.15 K using the Nosé–Hoover thermostat with a coupling constant of 1 ps. Pressure was maintained at 1 bar using a semi-isotropic Parrinello– Rahman barostat with a compressibility of 4.5 × 10^-5^ bar^-1^ and a coupling constant of 5 ps. For each mutant, three independent production runs were performed, with a new system generated from CHARMM-GUI for each replicate.

### Simulations Analysis

#### Hydrogen-Bonding Network Analysis

We used Bridge2(39) to visualize the hydrogen-bonding network between polar sidechains and/or bridging pore waters. We used the Bridge2 algorithm to detect hydrogen-bonding interactions between pore-facing residues and up to five bridging water molecules in the last 500 frames (100-200 ns) of each simulation. Results were filtered to exclude edges of the network with <10% occupancy and any residues outside the core (residues 6-20).

#### Water Density Plots and Hydrophobic Length Calculations

Water density analysis and plots were made using the Channel Annotation Package (CHAP)(40). The production files were reprocessed with GROMACS 2018 to generate a tpr file for CHAP processing. To generate the trajectory file, the coordinates of the simulation were saved every 200 ps. From these files, the water density for each frame along the z-axis of the pore could be extracted and plotted as shown in **Fig. 4G-J**. Average water density over the full length of the z-axis for the full simulation was then plotted along with the second derivative of the curve (*e*.*g*., **Supplementary Figure S6C-D**). Inflection points crossing the y axis of the second derivative were assigned as points of changing hydrophobicity and hydrophobic lengths of the pore were calculated as the distance between inflection points (**Supplementary Table S6**).

We calculated the water lifetime of molecules within the pore across the simulations. From the MDAnalysis results, all water coordinates for the whole simulation were extracted. For each frame, a pentagonal cylinder approximating the pore volume was defined using the Cα coordinates of residues 5 and 22 from all five chains. Water molecules within this geometric region were then identified and used for subsequent analysis. Water molecules were then tracked to determine the frames at which they entered and exited the cylinder, allowing calculation of their residence time within the pore (**Supplementary Figure S23**).

#### Sidechain Conformation and Dynamics Analysis

Additional analysis was performed using MDAnalysis to extract atom coordinates, dihedral angles, and hydrogen bond interactions for each 200 ps frame. 3D representations (*e*.*g*., **Fig. 4C-F**) of Gln sidechain orientation were generated by defining a vector from the Cα atom to the sidechain Nε2 atom for each pore-lining Gln residue and tracking its orientation over the course of the trajectory. Each plot contains 1,000 frames from a single production run. Time traces of the Gln vector for each residue in the helical bundle (*e*.*g*., **Fig. 6B**) were plotted as a function of simulation time. Histograms of the z-coordinate of the Gln vector (*e*.*g*., **Fig. 6A**) were generated by aggregating vector coordinates from all Gln residues across 1,000 frames from each of three independent production runs (n = 5 × 1,000 × 3 = 15,000).

Based on the histogram of Gln z-coordinates in **Fig. 6A**, Gln residues were classified into three separate states: ΔZ < 0.0 (downward facing), 0.0 < ΔZ < 2.4 (neutral facing), or ΔZ > 2.4 (upward facing). In each frame, all five pore-lining Gln residues were compared pairwise. A binary indicator was assigned to each pair (1 if both residues occupied the same state, 0 otherwise). This was computed for all Gln pairs across 1000 frames from each of three simulations. The values were then averaged to yield the pairwise state agreement between residues, defined as the fraction of frames in which a given pair occupies the same state. An agreement value of 1.0 indicates that two residues are always in the same state, whereas a value of 0.0 indicates that they never share the same state. The resulting agreement values are presented as a heatmap summarizing the relationships among the five Gln residues (*e*.*g*., **Fig. 6C**).

χ_2_ vs. χ_1_ plots (*e*.*g*., **Fig. 5E**) were generated using the dihedral angles extracted for all five Gln in the 1000-frame simulation for a single replicate. To build the background contour map, a database was curated with all PDBs annotated as a membrane protein from PDBTM(58), mpstruc(59), OPM(55), MemProtMD(60), with resolution <3.5Å. All *de novo* transmembrane proteins were manually added. The transmembrane regions were then filtered from each structure with PPM(55), resulting in a database of only the transmembrane regions from the structures. We then extracted the Gln sidechain dihedral angles for all instances of Gln in this database. From the χ_1_/χ_2_ values a 2D Gaussian Kernel Density Estimate was calculated and the 70^th^ and 90^th^ percentiles were shown as light and dark gray respectively. All dihedral angles extracted from simulation frames were plotted and clustered using DBSCAN clustering (**Supplementary Table 8**). For **Fig. 5**, we also extracted and plotted the Gln dihedral angles from our crystal structures (**Supplementary Table 9**).

### Peptide Synthesis

All peptides (including AM2-TM and mutants of LLLL) were synthesized using an automated Biotage Syro I parallel solid-phase peptide synthesizer. Peptides were synthesized on TentaGel S-RAM resin (CHEM-Impex) with a load of 0.23 mmol/g to produce a carboxamide C-terminus during cleavage. For AM2-TM only, the peptide was acetylated on the N-terminus using a mixture of 90:7:2 N,N-Dimethylformamide(DMF):N,N-Diisopropylethylamine:acetic anhydride and washed with DMF. All peptides were then washed with dichloromethane and cleaved from the resin using a solution of 95:2.5:2.5 trifluoroacetic acid (TFA):triisopropylsilane:water. Following cleavage, the samples were precipitated with cold diethyl ether, dissolved with 1,1,1,3,3,3-hexafluoroisopropanol, and then diluted by 50% with Solution A (99.9:0.1 water:TFA) for HPLC. HPLC purification of the peptides was performed on a C4 preparatory column (HiCHROM) from Solution A to Solution B (60:30:9.9:0.1 isopropanol:acetonitrile:water:TFA) with a 60-100% gradient of Solution B at a flow rate of 10 mL min^-1^. After HPLC, the peptides were lyophilized to dryness and then dissolved in ethanol for the final stock. Peptide identity and purity (**Supplementary Table 1**) were confirmed using MALDI-TOF spectrometry on a Bruker UltrafleXtreme with a-cyano-4-hydroxycinnamic acid (Sigma-Aldrich).

### SDS-PAGE Gel Electrophoresis

Samples for gel electrophoresis were prepared by lyophilizing enough of the stock solution to have 100 mg of each peptide. This was then dissolved in 25 mM Tris pH 7.5, 50 mM octyl-β-glucopyranoside (OG, ThermoScientific) to a final concentration of 2.5 mg mL^-1^ and then diluted 1:1 with 2x SDS loading dye. These samples were boiled for 10 minutes at 95°C and then loaded into the NuPAGE 12% Bis-Tris gel (Invitrogen) in MES buffer. The final load for each protein was 4 ug. The gel was then run at 200 mV for 30 minutes, stained with Coomassie dye, and then imaged.

### Proton Flux Assay

The protocol for the preparation of the proteoliposomes was adapted from methods described in Moffat *et al*.*(37)*, Ma *et al*.*(38)*, and Kratochvil *et al*.*(36)*. We describe the exact methods used in our study below.

Lipids were prepared in stocks of 1-palmitoyl-2-oleoyl-glycero-3-phosphocholine (16:0-18:1 PC, POPC, Avanti) in ethanol at 100 mg/mL and 1-palmitoyl-2-oleoyl-sn-glycero-3-phospho-(1’-rac-glycerol) sodium salt (16:0-18:1 PG, POPG, Avanti) in chloroform at 25 mg/mL. 2 mL of 3:1 PC:PG lipid stock solutions were prepared by adding 150 mL of POPC ethanol stock and 200 mL of POPG chloroform stock to a scintillation vial. Drying under an air stream for 30 minutes produced a film of lipids, which was lyophilized overnight to remove all organic solvents. The lipid film was resuspended in 2 mL of K^+^ buffer (50 mM potassium sulfate, 30 mM potassium phosphate, pH 7.5) with 500 uM pyranine (HPTS, SigmaAldrich). This lipid stock solution was then tip-sonicated in 1 mL batches for 10 min at 20% power using 5 s on/off cycles. The sonicated solutions were then recombined prior to three freeze/thaw cycles. After the third cycle, lipids were extruded in 1 mL aliquots through a 200 nm filter with gas-tight Hamilton syringes at room temperature. A stock of 20% Triton X100 (TX100, ThermoFisher) in water was then used to reach a final concentration of 0.1% TX100 in the solution. Liposomes were then left on a rotisserie at room temperature for 30 minutes to equilibrate. At this stage, the liposomes were uniform and ready for peptide addition.

Peptide stocks for liposomal assays were prepared by lyophilizing enough ethanol stock to result in 100 mg of peptide. These were then dissolved in 1% TX100 in water to produce a final stock concentration of 1.77 mM peptide. AM2-TM was dissolved in 2% OG in water at the same final peptide concentration. The liposome with TX100 solution was then aliquoted into 100 mL aliquots and diluted at a 1:5 ratio with 400 uL of K^+^ buffer with 500 uM HPTS and 0.1% TX100. Aliquots for AM2-TM were not diluted. 8 mL of peptide stock solution was added to 4 aliquots for each mutant to generate four biological replicates of peptide:lipid molar ratio of 1:100. For empty liposome samples, 8 mL of 1% TX100 was added. Samples were left on the rotisserie for 30 minutes at room temperature. To remove detergent, an Amberlite XAD-2 (Sigma-Aldrich) biobeads slurry was prepared in K^+^ buffer. A total of 120 µL (∼60 mg) was added to each aliquot and incubated on a rotisserie for 1 h at 4 °C. This step was repeated once, followed by the addition of 240 µL (∼120 mg) and overnight incubation at 4 °C. In total, 480 µL (∼240 mg) of biobeads were added per sample. Samples were then spun down in a benchtop centrifuge at 20,000x*g* for 15 minutes to pellet the biobeads. 500 mL of supernatant was removed and then spun down in an ultracentrifuge at 120,000x*g* to pellet the proteoliposomes. The supernatant was removed, and the pellet was resuspended in 100 μL of K^+^ buffer to generate the proteoliposome stock solution.

The proton flux assay was run on a Biotek Synergy H1 Plate Reader. Assay samples were prepared by taking 13 mL of the proteoliposome stock solution and adding 637 mL of Na^+^ buffer (50 mM sodium sulfate, 30 mM sodium phosphate, pH 7.5) with 10 mM p-xylene-bis-pyridinium bromide (DPX, Invitrogen) to quench any extraliposomal HPTS fluorescence. Samples were left to incubate at room temperature in the dark for 25 minutes. Working solutions of the potassium ionophore valinomycin (SigmaAldrich) and the protonophore carbonyl cyanide m-chlorophenyl hydrazone (CCCP, Cayman Chemical Company) were prepared at 5 mM in Na^+^ buffer. Technical triplicates were performed by dispensing 190 µL of assay sample into three wells of a black 96-well plate. Fluorescence was recorded at an emission wavelength of 518 nm with excitation at 417 nm (pH-independent) and 460 nm (pH-dependent). Data are reported as the ratiometric signal at 460 nm to 417 nm excitation. Fluorescence was monitored every 10 s for 2 min prior to the addition of 10 µL valinomycin working solution to each well. Measurements continued at 10 s intervals for 10 min, followed by the addition of 10 µL CCCP and an additional 5 min of data collection. Each of the reported samples included eight biological replicates, each performed in technical triplicate (24 total technical measurements).

A calibration curve (**Supplementary Figure S1**) was generated by preparing HPTS in K^+^ buffer over a pH range of 3–9. The fluorescence ratio (F_460nm_/F_417nm_) was fitted to a sigmoidal function to correlate ratiometric signal with pH according to the following equation:

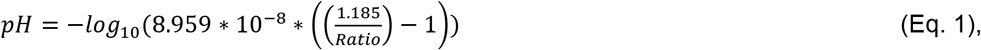

where Ratio represents the ratio of fluorescent signal between the deprotonated dye (*λ*_emission_ = 460 nm, *λ*_excitation_ = 515 nm) and the isosbestic point (*λ*_emission_ = 417 nm, *λ*_excitation_ = 515 nm). These ratio values measured with the plate readers were then converted to pH.

Finally, readings were converted to total proton concentration by also factoring in the buffer capacity of phosphate buffer according to the following equation:

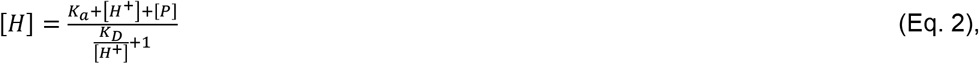

where K_a_ is the acid dissociation constant of phosphate (6.31*10^-8^) for the pH range observed (pH=3-9), [H+] is the free proton concentration or the -log(pH) obtained from fluorescence signal, and [P] is the total concentration of phosphate (0.03 M). Total proton concentration curves obtained from the points within the first 60 seconds after valinomycin was added were fitted with a linear model, and the slope or rates were plotted in **Fig. 3C-E**.

### Lipidic Cubic Phase Crystallography

Methods for lipidic cubic phase (LCP) crystallography were adapted from Caffrey and Cherezov(61, 62) and Kratochvil *et al*.*(36)*. Briefly, peptide-lipid sample was first prepared by adding 750 μg of peptide from the peptide ethanol stock to 30 mg of monoolein (Hampton Research) and mixed until clear. Ethanol was evaporated under an air stream, and then samples were lyophilized overnight to remove any residual solvent. Samples were melted in a 56°C bead bath and then transferred to a gas-tight Hamilton 250 mL syringe. Based on the mass of the melted peptide–lipid mixture, an aqueous phase containing 50 mM OG was added at two-thirds of the mixture mass to achieve a final composition of 40% (w/w) aqueous phase. The mixture was extruded 200 times at room temperature using an LCP coupler fitted with a 100 µL syringe.

Crystal trays were set using the Formulatrix NT8 at room temperature on 96-well Laminex plastic sandwich plates (Molecular Dimensions). Wells were prepared with 50 nL of the extruded LCP and 1 uL of precipitant solution. Plates were monitored using the Formulatrix RI-1 Imager. Conditions that produced crystals with high-quality diffraction are described in **Supplementary Table S10**. Crystals were looped, cryoprotected in 30% PEG 400 if necessary, and then frozen and stored in liquid nitrogen.

Diffraction data was collected at the Stanford Synchrotron Radiation Lightsource on beamlines 12-1 and 12-2 with wavelengths of 1.14Å and 0.98Å respectively. The data was processed using DIALS(63). All further steps of structure solving were performed within the CCP4 suite(64). The data was merged and reduced with AIMLESS(65), and molecular replacement was performed with Phaser(66) using the structure for LLLL (PDB 6MCT) as the initial model. The structure was refined with Refmac5(67) and the model was built using COOT(68). PDB-REDO(69) was also used to optimize refinement. Final model statistics are reported in **Supplementary Table S11**.

## Acknowledgments

We thank E.S.B., K.M.J., and A.N.F. for helping with the statistical analysis methods presented in the publication. We also thank those involved with the CCP4/APS School in Macromolecular Crystallography 2024 for help with X-ray crystallography data processing and structure refinement. N.P.J. was supported by the Molecular and Cellular Biophysics NIGMS T32 training grant T32GM148376. V.T.S. was supported by the Chemical Biology Interface NIGMS T32 training grant T32GM135122. H.T.K. acknowledges support from the NIH NIGMS R00GM138753. Use of the Stanford Synchrotron Radiation Lightsource, SLAC National Accelerator Laboratory, is supported by the U.S. Department of Energy, Office of Science, Office of Basic Energy Sciences under Contract No. DE-AC02-76SF00515. The SSRL Structural Molecular Biology Program is supported by the DOE Office of Biological and Environmental Research, and by the National Institutes of Health, National Institute of General Medical Sciences (P30GM133894). The contents of this publication are solely the responsibility of the authors and do not necessarily represent the official views of NIGMS or NIH.

## References

1. P. Mitchell, Coupling of Phosphorylation to Electron and Hydrogen Transfer by a Chemi-Osmotic type of Mechanism. Nature 191, 144–148 (1961).

2. D. G. Nicholls, Mitochondrial ion circuits. Essays in Biochemistry 47, 25–35 (2010).

3. T. Nishi, M. Forgac, The vacuolar (H+)-ATPases — nature’s most versatile proton pumps. Nat Rev Mol Cell Biol 3, 94–103 (2002).

4. G. H. Diering, M. Numata, Endosomal pH in neuronal signaling and synaptic transmission: role of Na+/H+ exchanger NHE5. Front. Physiol. 4 (2014).

5. Y. Moriyama, M. Futai, H+-ATPase, a primary pump for accumulation of neurotransmitters, is a major constituent of brain synaptic vesicles. Biochemical and Biophysical Research Communications 173, 443–448 (1990).

6. N. Agmon, The Grotthuss mechanism. Chemical Physics Letters 244, 456–462 (1995).

7. P. B. Calio, C. Li, G. A. Voth, Resolving the Structural Debate for the Hydrated Excess Proton in Water. J. Am. Chem. Soc. 143, 18672–18683 (2021).

8. C. Knight, G. A. Voth, The Curious Case of the Hydrated Proton. Acc. Chem. Res. 45, 101– 109 (2012).

9. T. E. Decoursey, Voltage-Gated Proton Channels and Other Proton Transfer Pathways. Physiological Reviews 83, 475–579 (2003).

10. A.-N. Bondar, “Mechanisms of long-distance allosteric couplings in proton-binding membrane transporters” in Advances in Protein Chemistry and Structural Biology, (Elsevier, 2022), pp. 199–239.

11. R. H. Tunuguntla, F. I. Allen, K. Kim, A. Belliveau, A. Noy, Ultrafast proton transport in sub-1-nm diameter carbon nanotube porins. Nature Nanotech 11, 639–644 (2016).

12. J. Geng, et al., Stochastic transport through carbon nanotubes in lipid bilayers and live cell membranes. Nature 514, 612–615 (2014).

13. K. Otake, et al., Confined water-mediated high proton conduction in hydrophobic channel of a synthetic nanotube. Nat Commun 11, 843 (2020).

14. L. Xu, et al., Rational Tuning the Proton Conductivity and Stability of Hydrogen-Bonded Organic Frameworks. Inorg. Chem. 63, 17747–17754 (2024).

15. D. G. Levitt, S. R. Elias, J. M. Hautman, Number of water molecules coupled to the transport of sodium, potassium and hydrogen ions via gramicidin, nonactin or valinomycin. Biochimica et Biophysica Acta (BBA) - Biomembranes 512, 436–451 (1978).

16. P. A. Rosenberg, A. Finkelstein, Interaction of ions and water in gramicidin A channels: streaming potentials across lipid bilayer membranes. The Journal of general physiology 72, 327– 340 (1978).

17. J. R. Schnell, J. J. Chou, Structure and mechanism of the M2 proton channel of influenza A virus. Nature 451, 591–595 (2008).

18. S. D. Cady, et al., Structure of the amantadine binding site of influenza M2 proton channels in lipid bilayers. Nature 463, 689–692 (2010).

19. M. Wikström, K. Krab, V. Sharma, Oxygen Activation and Energy Conservation by Cytochrome c Oxidase. Chem. Rev. 118, 2469–2490 (2018).

20. M. Tashiro, A. A. Stuchebrukhov, Thermodynamic Properties of Internal Water Molecules in the Hydrophobic Cavity around the Catalytic Center of Cytochrome c Oxidase. J. Phys. Chem. B 109, 1015–1022 (2005).

21. P. Goyal, J. Lu, S. Yang, M. R. Gunner, Q. Cui, Changing hydration level in an internal cavity modulates the proton affinity of a key glutamate in cytochrome c oxidase. Proc. Natl. Acad. Sci. U.S.A. 110, 18886–18891 (2013).

22. I. S. Ramsey, et al., An aqueous H+ permeation pathway in the voltage-gated proton channel Hv1. Nat Struct Mol Biol 17, 869–875 (2010).

23. H. Jain, M. Lazaratos, P. Pohl, A.-N. Bondar, Fluctuating hydrogen-bond network of the Hv1 ion channel. Comput Struct Biotechnol J 27, 4403–4417 (2025).

24. C. Li, G. A. Voth, A quantitative paradigm for water-assisted proton transport through proteins and other confined spaces. Proc. Natl. Acad. Sci. U.S.A. 118, e2113141118 (2021).

25. Y. Peng, J. M. J. Swanson, S. Kang, R. Zhou, G. A. Voth, Hydrated Excess Protons Can Create Their Own Water Wires. J. Phys. Chem. B 119, 9212–9218 (2015).

26. N. H. Joh, et al., De novo design of a transmembrane Zn2+-transporting four-helix bundle. Science (2014). 10.1126/science.1261172.

27. A. Vorobieva, et al., De novo design of transmembrane β barrels. Science (2021). 10.1126/science.abc8182.

28. A. J. Scott, et al., Constructing ion channels from water-soluble α-helical barrels. Nat. Chem. 13, 643–650 (2021).

29. S. Berhanu, et al., Sculpting conducting nanopore size and shape through de novo protein design. Science 385, 282–288 (2024).

30. Y. Liu, et al., Bottom-up design of Ca2+ channels from defined selectivity filter geometry. Nature 648, 468–476 (2025).

31. C. Zhou, et al., De novo designed voltage-gated anion channels suppress neuron firing. Cell 188, 7495–7511.e21 (2025).

32. J. D. Lear, Z. R. Wasserman, W. F. DeGrado, Synthetic Amphiphilic Peptide Models for Protein Ion Channels. Science 240, 1177–1181 (1988).

33. Y. Wu, G. A. Voth, A Computer Simulation Study of the Hydrated Proton in a Synthetic Proton Channel. Biophysical Journal 85, 864–875 (2003).

34. Y. Wu, B. Ilan, G. A. Voth, Charge Delocalization in Proton Channels, II: The Synthetic LS2 Channel and Proton Selectivity. Biophysical Journal 92, 61–69 (2007).

35. M. Mravic, et al., Packing of apolar side chains enables accurate design of highly stable membrane proteins. Science 363, 1418–1423 (2019).

36. H. T. Kratochvil, et al., Transient water wires mediate selective proton transport in designed channel proteins. Nat. Chem. 15, 1012–1021 (2023).

37. J. C. Moffat, et al., Proton Transport through Influenza A Virus M2 Protein Reconstituted in Vesicles. Biophysical Journal 94, 434–445 (2008).

38. C. Ma, et al., Identification of the functional core of the influenza A virus A/M2 proton-selective ion channel. Proc. Natl. Acad. Sci. U.S.A. 106, 12283–12288 (2009).

39. M. Siemers, A.-N. Bondar, Interactive Interface for Graph-Based Analyses of Dynamic H-Bond Networks: Application to Spike Protein S. J. Chem. Inf. Model. 61, 2998–3014 (2021).

40. G. Klesse, S. Rao, M. S. P. Sansom, S. J. Tucker, CHAP: A Versatile Tool for the Structural and Functional Annotation of Ion Channel Pores. Journal of Molecular Biology 431, 3353–3365 (2019).

41. Y. Pankratova, M. Hong, Side Chain Structures of the Proton-Selective Histidine and Gating Tryptophan in Influenza BM2 Reveal Both Conservation and Variation of the Proton Conduction Mechanism. Biochemistry 64, 3081–3092 (2025).

42. S. K. Nair, D. W. Christianson, Unexpected pH-dependent conformation of His-64, the proton shuttle of carbonic anhydrase II. J. Am. Chem. Soc. 113, 9455–9458 (1991).

43. A. B. Weinglass, I. N. Smirnova, H. R. Kaback, Engineering Conformational Flexibility in the Lactose Permease of Escherichia coli : Use of Glycine-Scanning Mutagenesis To Rescue Mutant Glu325→Asp. Biochemistry 40, 769–776 (2001).

44. S. Roy, et al., Mechanism of Side Chain-Controlled Proton Conductivity in Bioinspired Peptidic Nanostructures. J. Phys. Chem. B 125, 12741–12752 (2021).

45. S. Taraphder, G. Hummer, Protein Side-Chain Motion and Hydration in Proton-Transfer Pathways. Results for Cytochrome P450cam. J. Am. Chem. Soc. 125, 3931–3940 (2003).

46. T. J. Harpole, C. Grosman, Side-chain conformation at the selectivity filter shapes the permeation free-energy landscape of an ion channel. Proc. Natl. Acad. Sci. U.S.A. 111 (2014).

47. Z. Li, et al., Asymmetric gating of a homopentameric ion channel GLIC revealed by cryo-EM. Proc. Natl. Acad. Sci. U.S.A. 122, e2512811122 (2025).

48. S. Minniberger, S. Abdolvand, S. Braunbeck, H. Sun, A. J. R. Plested, Asymmetry and Ion Selectivity Properties of Bacterial Channel NaK Mutants Derived from Ionotropic Glutamate Receptors. Journal of Molecular Biology 435, 167970 (2023).

49. A. Di Luca, A. P. Gamiz-Hernandez, V. R. I. Kaila, Symmetry-related proton transfer pathways in respiratory complex I. Proc. Natl. Acad. Sci. U.S.A. 114 (2017).

50. X. Hou, H. Zhang, L. Jiang, Building Bio-Inspired Artificial Functional Nanochannels: From Symmetric to Asymmetric Modification. Angew Chem Int Ed 51, 5296–5307 (2012).

51. H. Zhang, Y. Tian, L. Jiang, From symmetric to asymmetric design of bio-inspired smart single nanochannels. Chem. Commun. 49, 10048 (2013).

52. S. Jo, T. Kim, W. Im, Automated Builder and Database of Protein/Membrane Complexes for Molecular Dynamics Simulations. PLoS ONE 2, e880 (2007).

53. S. Jo, T. Kim, V. G. Iyer, W. Im, CHARMM-GUI: A web-based graphical user interface for CHARMM. J Comput Chem 29, 1859–1865 (2008).

54. E. L. Wu, et al., CHARMM-GUI Membrane Builder toward realistic biological membrane simulations. J. Comput. Chem. 35, 1997–2004 (2014).

55. M. A. Lomize, I. D. Pogozheva, H. Joo, H. I. Mosberg, A. L. Lomize, OPM database and PPM web server: resources for positioning of proteins in membranes. Nucleic Acids Research 40, D370–D376 (2012).

56. J. Huang, A. D. MacKerell, CHARMM36 all-atom additive protein force field: Validation based on comparison to NMR data. J. Comput. Chem. 34, 2135–2145 (2013).

57. D. Van Der Spoel, et al., GROMACS: Fast, flexible, and free. J Comput Chem 26, 1701– 1718 (2005).

58. D. Kozma, I. Simon, G. E. Tusnády, PDBTM: Protein Data Bank of transmembrane proteins after 8 years. Nucleic Acids Research 41, D524–D529 (2012).

59. S. H. White, Biophysical dissection of membrane proteins. Nature 459, 344–346 (2009).

60. T. D. Newport, M. S. P. Sansom, P. J. Stansfeld, The MemProtMD database: a resource for membrane-embedded protein structures and their lipid interactions. Nucleic Acids Research 47, D390–D397 (2019).

61. M. Caffrey, V. Cherezov, Crystallizing membrane proteins using lipidic mesophases. Nat Protoc 4, 706–731 (2009).

62. M. Caffrey, Crystallizing Membrane Proteins for Structure Determination: Use of Lipidic Mesophases. Annu. Rev. Biophys. 38, 29–51 (2009).

63. G. Winter, et al., DIALS : implementation and evaluation of a new integration package. Acta Crystallogr D Struct Biol 74, 85–97 (2018).

64. M. D. Winn, et al., Overview of the CCP 4 suite and current developments. Acta Crystallogr D Biol Crystallogr 67, 235–242 (2011).

65. P. R. Evans, G. N. Murshudov, How good are my data and what is the resolution? Acta Crystallogr D Biol Crystallogr 69, 1204–1214 (2013).

66. A. J. McCoy, et al., Phaser crystallographic software. J Appl Crystallogr 40, 658–674 (2007).

67. G. N. Murshudov, et al., REFMAC 5 for the refinement of macromolecular crystal structures. Acta Crystallogr D Biol Crystallogr 67, 355–367 (2011).

68. P. Emsley, B. Lohkamp, W. G. Scott, K. Cowtan, Features and development of Coot. Acta Crystallogr D Biol Crystallogr 66, 486–501 (2010).

69. R. P. Joosten, F. Long, G. N. Murshudov, A. Perrakis, The PDB_REDO server for macromolecular structure model optimization. IUCrJ 1, 213–220 (2014).

