## Supplementary material for "Engineered Channel Asymmetry Extends Hydrogen-Bonding Networks for Proton Conduction": SI Appendix

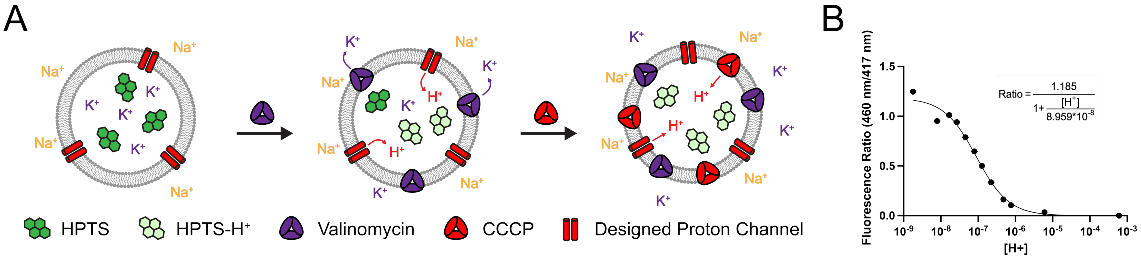

**Supplementary Figure 1.** (A) Cartoon schematic of liposomal assay including valinomycin and CCCP addition steps. The CCCP step is included to test for liposome leakiness. (B) Calibration curve for HPTS over a range of pH 3-9. HPTS samples were made at a concentration of ~10 uM in 50 mM potassium sulfate, 30 mM potassium phosphate, pH 7.5. Fluorescence ratio (460 nm/417 nm) were fitted with a sigmoidal curve and plotted to correlate fluorescence signal with pH. Each point contains n=3 measurements with SEM error bars. The R-squared value for the fit is 0.9861.

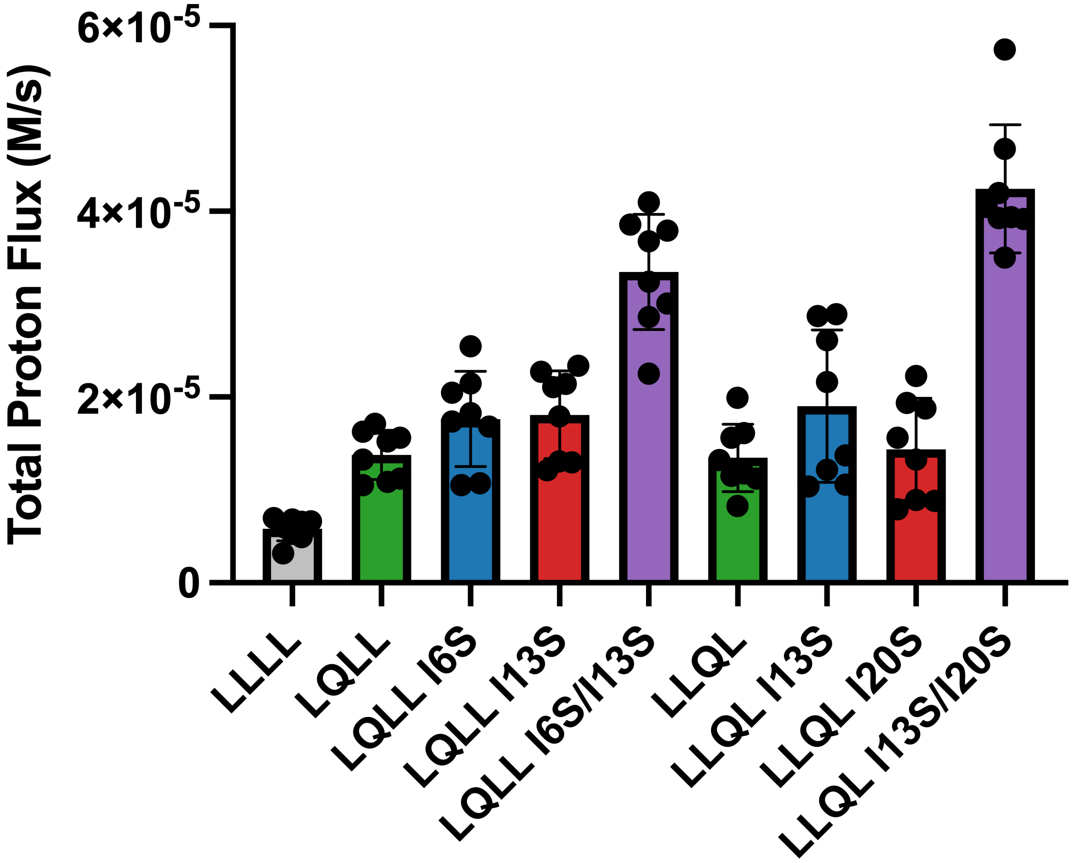

**Supplementary Figure 2. Initial flux rates for all mutants tested as well as the control bundle LLLL.** All variants were compared to LLLL with a one-way ANOVA and post-hoc Dunnett's test. All variants exhibit significantly higher proton conductivity than the completely hydrophobic pore LLLL (**Supplemental Table 3**).

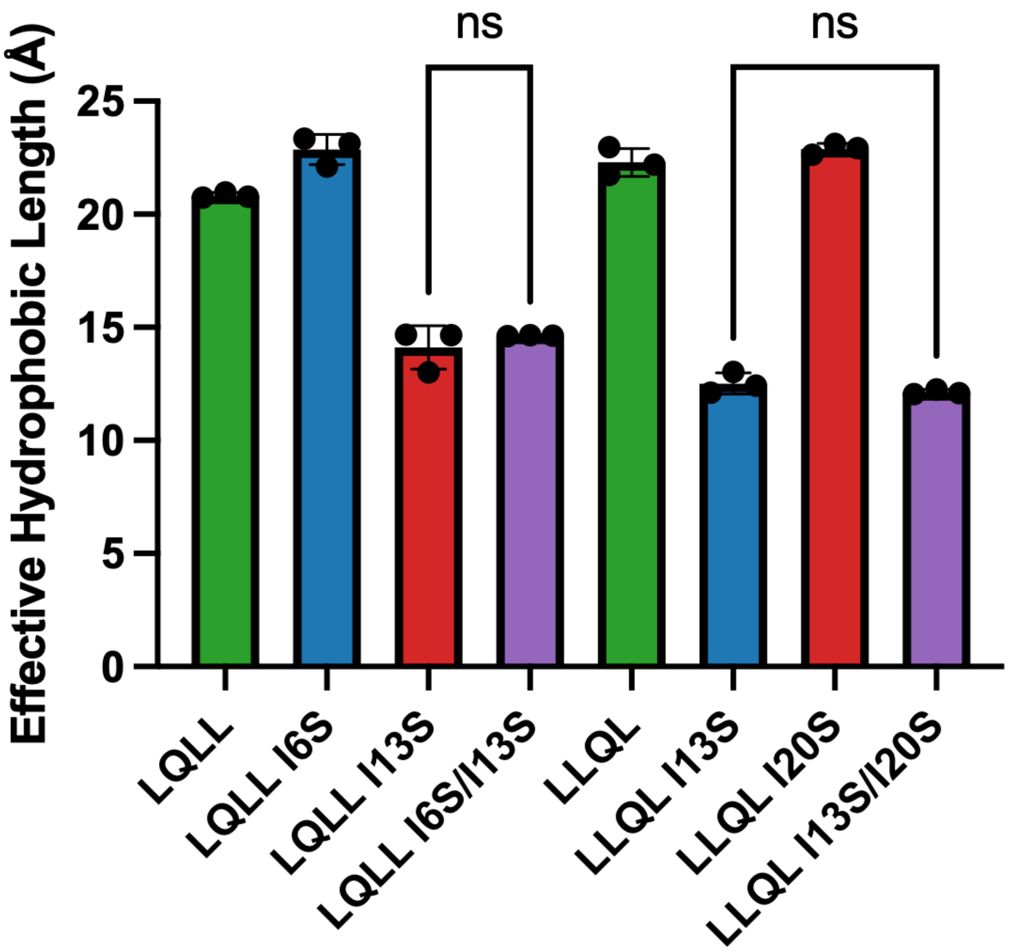

**Supplementary Figure 3. Effective hydrophobic length for all mutants based on average water density.** Bars represent the mean value across n=3 simulations with error bars showing SEM. To compare significance amongst mutants with shorter hydrophobic lengths, a one-way ANOVA and post-hoc Tukey test was performed across all mutants. Within families for mutants with lower hydrophobic lengths, there was no significant difference (**Supplemental Tables 12 and 13**). Results are also shown in **Supplemental Table 6**.

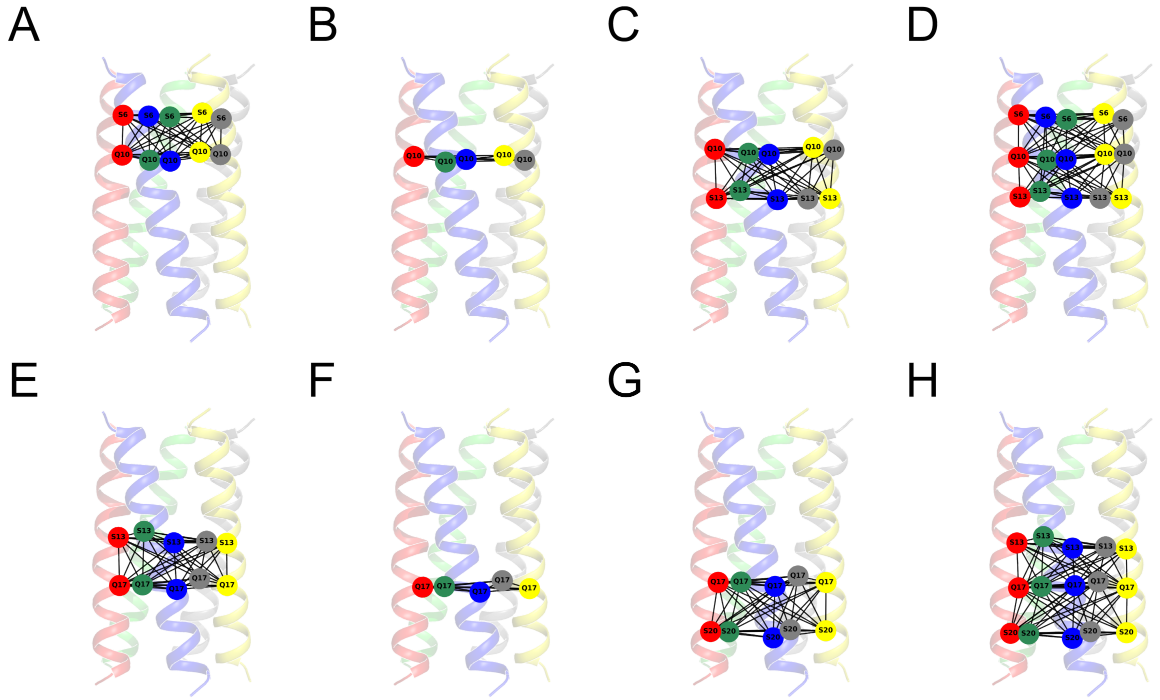

**Supplementary Figure 4. Bridge2 hydrogen-bonding network analysis.** For each mutant (A) LQLL I6S, (B) LQLL, (C) LQLL I13S, (D) LQLL I6S/I13S, (E) LLQL I13S, (F) LLQL, (G) LLQL I20S, and (H) LLQL I13S/I20S the hydrogen-bonding network between polar residues within the pore is shown. All networks are overlaid over the PDB structure. The structure and the nodes are colored by chain. Nodes represent polar residues within the pore. Edges represent hydrogen-bonding networks connecting two residues, containing at most five water molecules. The displayed edges have been filtered to require an occupancy greater than 10% of the 1000 frames in a single simulation production.

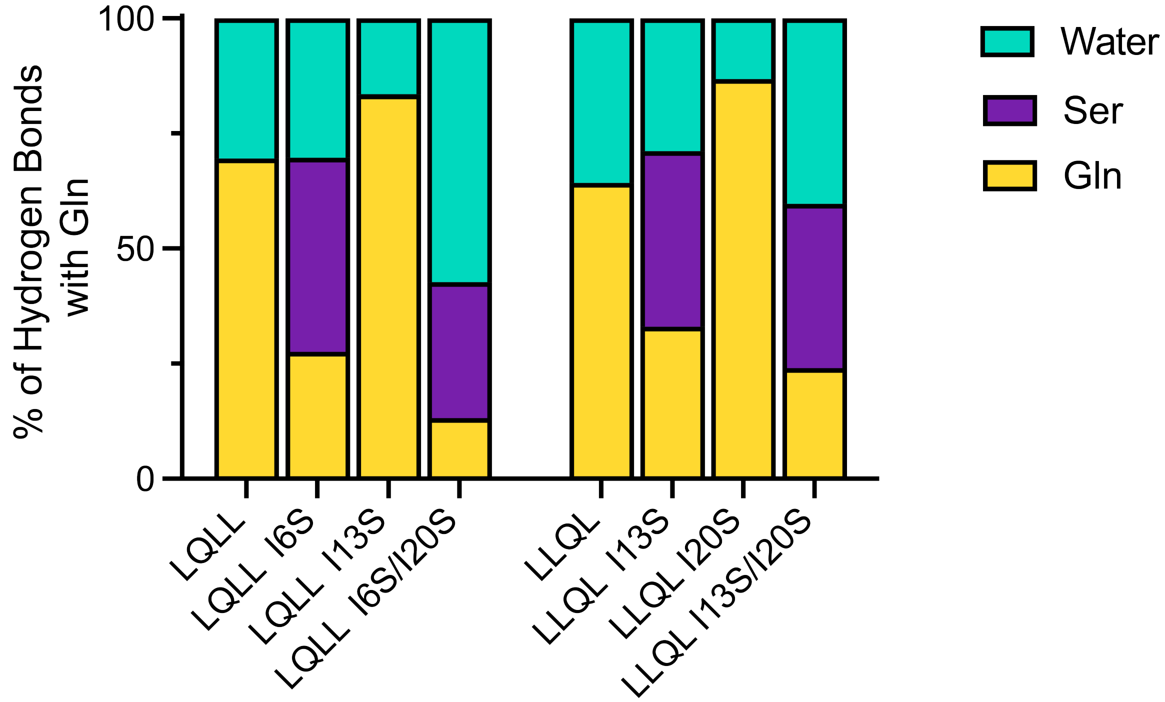

**Supplementary Figure 5. Hydrogen bonding partners with Gln10 or Gln17 throughout the MD simulations.** Bars represent the percent of total hydrogen bonds between Gln and its respective partners. Hydrogen bonds are totaled across all three simulations for each mutant. Results are also included in **Supplementary Table 14**.

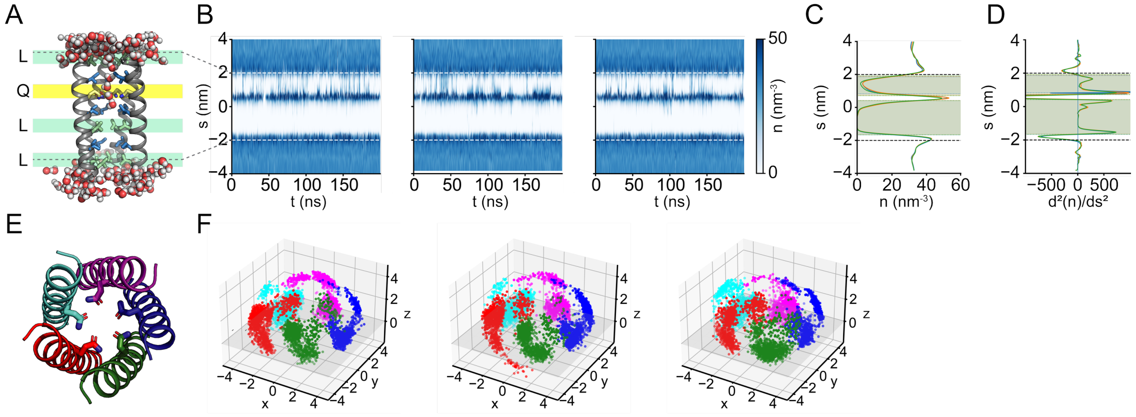

**Supplementary Figure 6. All MD simulation replicates for LQLL.** (A) Representative snapshot of LQLL. Waters within 3 Å of the protein are shown, and pore-facing residues are colored as in **Fig. 2**. (B) Water density profiles for all three production runs (1-3, left to right). Dashed lines represent the protein boundaries within the simulation. (C) Average water density line plots are shown for each production (1=blue, 2=orange, 3=green). (D) Second derivative of the average water density plot from (C). Shaded hydrophobic regions were determined by points where there is an intercept at x=0 within the protein boundaries (dashed). (E) Model pentamer with each of its five helices colored to denote the separate chains. (F) For all three production runs (1-3, left to right), $\vec{v}_{Gln}$ is plotted for the five Gln10 residues over the course of the simulation. The Cα plane is denoted in gray for (x, y, z = 0).

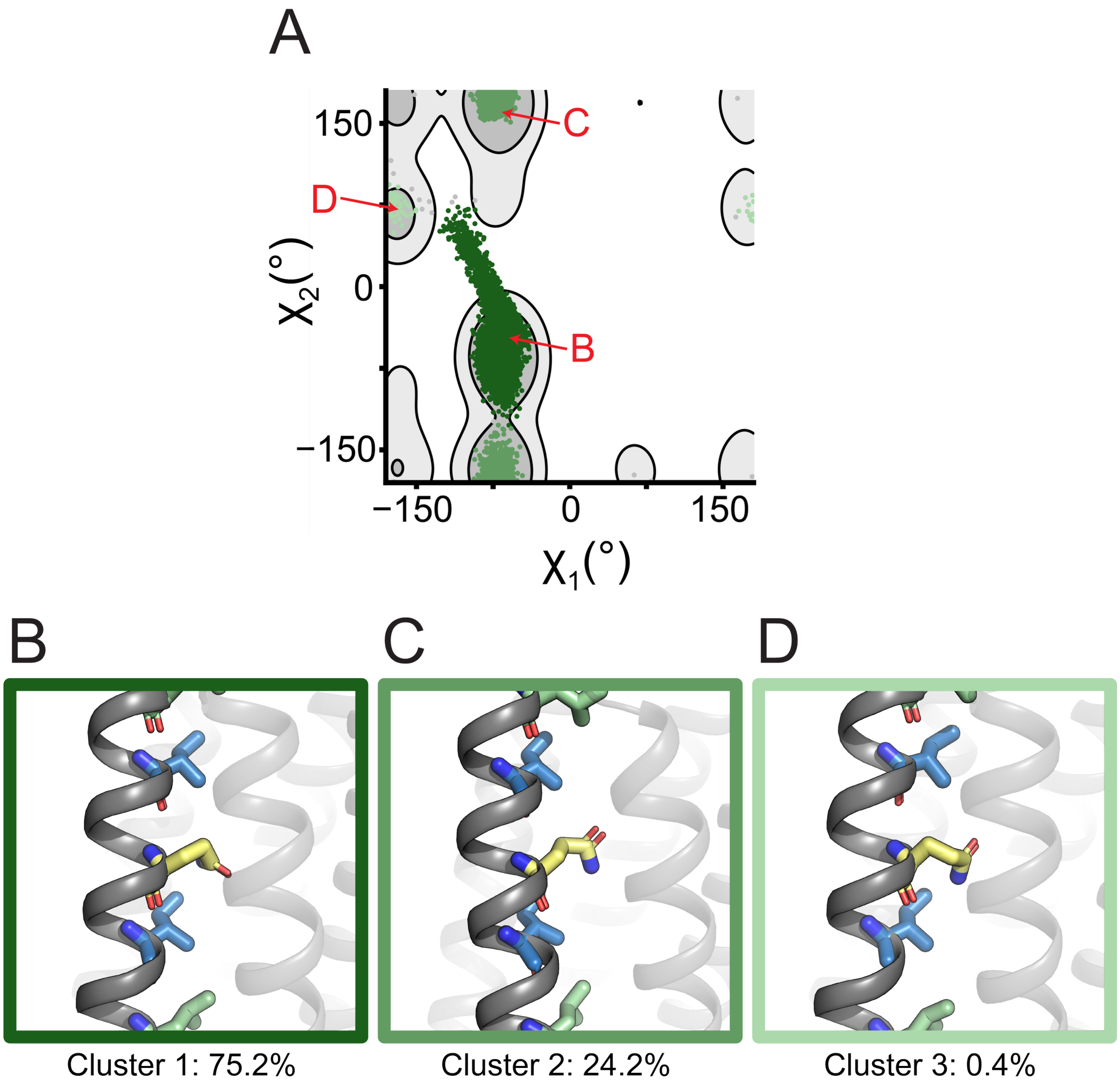

**Supplementary Figure 7. $\chi_1$/$\chi_2$ rotamer analysis of LQLL MD simulations.** (A) Plot of $\chi_1$/$\chi_2$ dihedral angles from Gln sidechains sampled from MD simulations for each of the mutants. These are overlaid on a contour map of Gln rotameric states sampled from a curated structural database of transmembrane proteins, where lighter gray contours denote the top 70% and darker gray contours denote the top 90%. Simulation points are colored by cluster with more prevalent clusters represented in darker colors. (B-D) Representative snapshots from each cluster are shown with pore-facing residues colored as in **Fig. 2**. Boxes are colored by cluster and labeled on the graph (A).

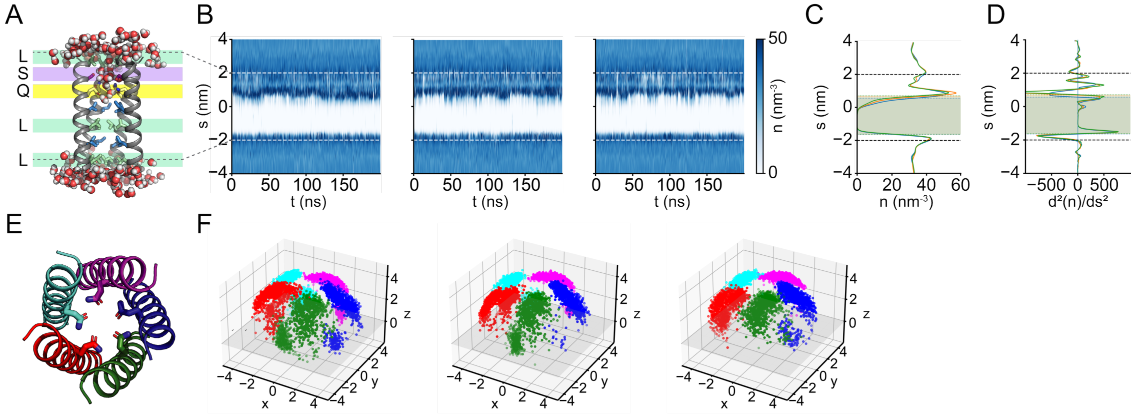

**Supplementary Figure 8. All MD simulation replicates for LQLL I6S.** (A) Representative snapshot of LQLL I6S. Waters within 3 Å of the protein are shown, and pore-facing residues are colored as in **Fig. 2**. (B) Water density profiles for all three production runs (1-3, left to right). Dashed lines represent the protein boundaries within the simulation. (C) Average water density line plots are shown for each production (1=blue, 2=orange, 3=green). (D) Second derivative of the average water density plot from (C). Shaded hydrophobic regions were determined by points where there is an intercept at x=0 within the protein boundaries (dashed). (E) Model pentamer with each of its five helices colored to denote the separate chains. (F) For all three production runs (1-3, left to right), $\vec{v}_{Gln}$ is plotted for the five Gln10 residues over the course of the simulation. The Cα plane is denoted in gray for (x, y, z = 0).

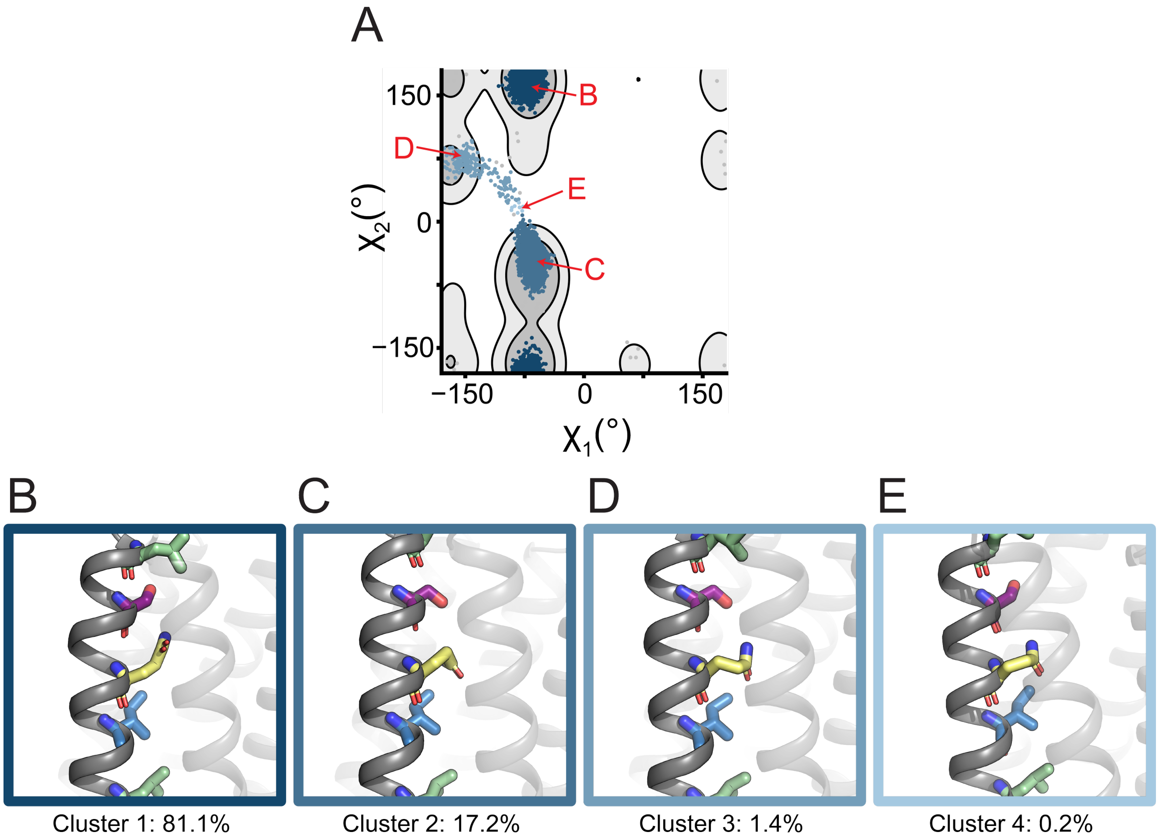

**Supplementary Figure 9. $\chi_1/\chi_2$ rotamer analysis of LQLL I6S MD simulations.** (A) Plot of $\chi_1/\chi_2$ dihedral angles from Gln sidechains sampled from MD simulations for each of the mutants. These are overlaid on a contour map of Gln rotameric states sampled from a curated structural database of transmembrane proteins, where lighter gray contours denote the top 70% and darker gray contours denote the top 90%. Simulation points are colored by cluster with more prevalent clusters represented in darker colors. (B-D) Representative snapshots from each cluster are shown with pore-facing residues colored as in **Fig. 2**. Boxes are colored by cluster and labeled on the graph (A).

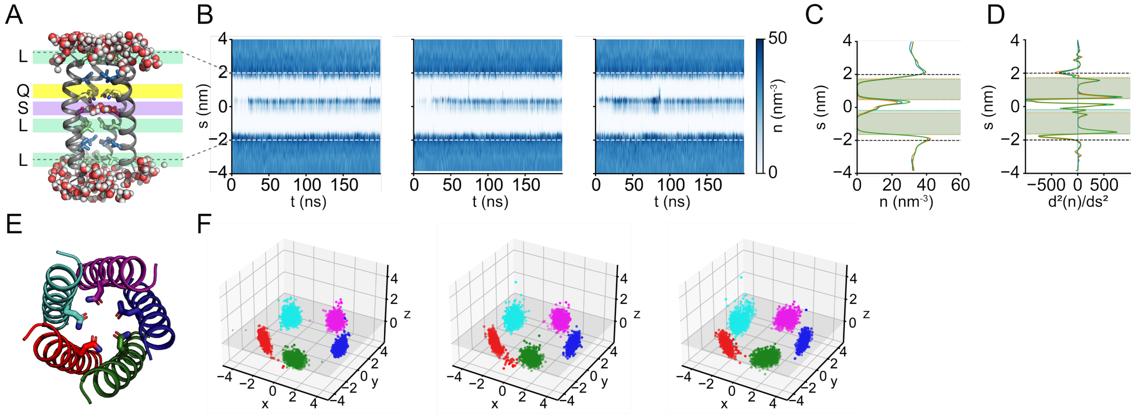

**Supplementary Figure 10. All MD simulation replicates for LQLL I13S.** (A) Representative snapshot of LQLL I13S. Waters within 3 Å of the protein are shown, and pore-facing residues are colored as in **Fig. 2**. (B) Water density profiles for all three production runs (1-3, left to right). Dashed lines represent the protein boundaries within the simulation. (C) Average water density line plots are shown for each production (1=blue, 2=orange, 3=green). (D) Second derivative of the average water density plot from (C). Shaded hydrophobic regions were determined by points where there is an intercept at x=0 within the protein boundaries (dashed). (E) Model pentamer with each of its five helices colored to denote the separate chains. (F) For all three production runs (1-3, left to right), $\vec{v}_{Gln}$ is plotted for the five Gln10 residues over the course of the simulation. The Cα plane is denoted in gray for (x, y, z = 0).

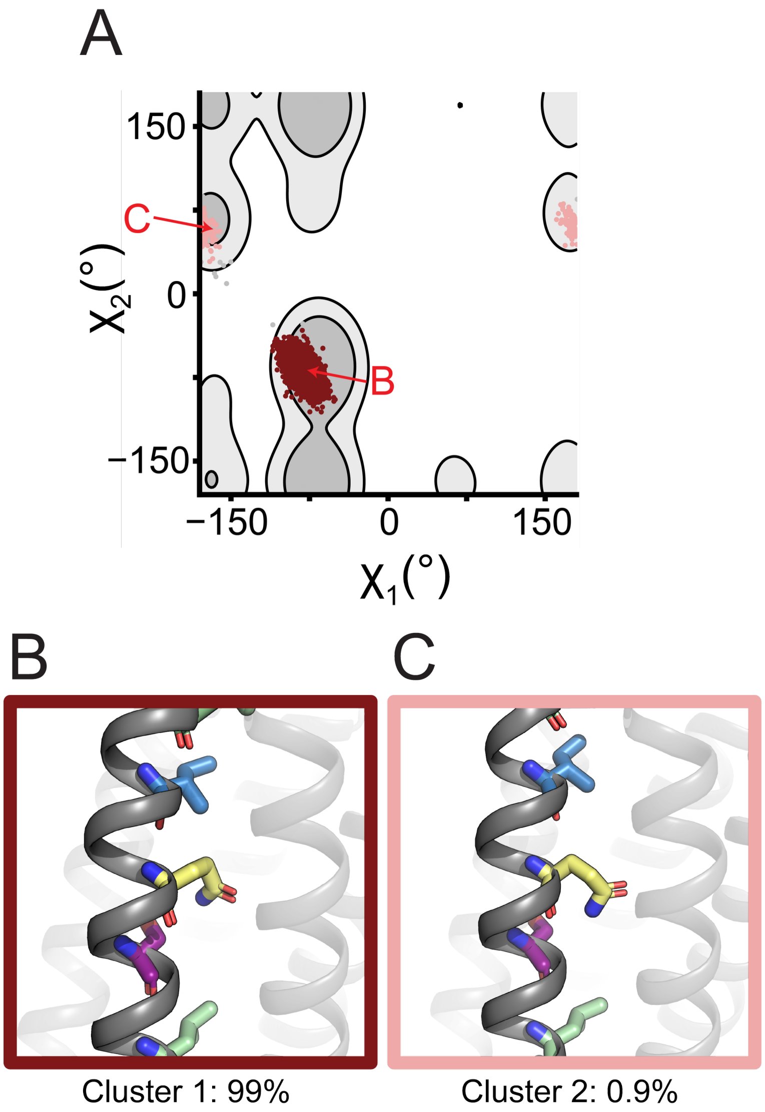

**Supplementary Figure 11. $\chi_1/\chi_2$ rotamer analysis of LQLL I13S MD simulations.** (A) Plot of $\chi_1/\chi_2$ dihedral angles from Gln sidechains sampled from MD simulations for each of the mutants. These are overlaid on a contour map of Gln rotameric states sampled from a curated structural database of transmembrane proteins, where lighter gray contours denote the top 70% and darker gray contours denote the top 90%. Simulation points are colored by cluster with more prevalent clusters represented in darker colors. (B-D) Representative snapshots from each cluster are shown with pore-facing residues colored as in **Fig. 2**. Boxes are colored by cluster and labeled on the graph (A).

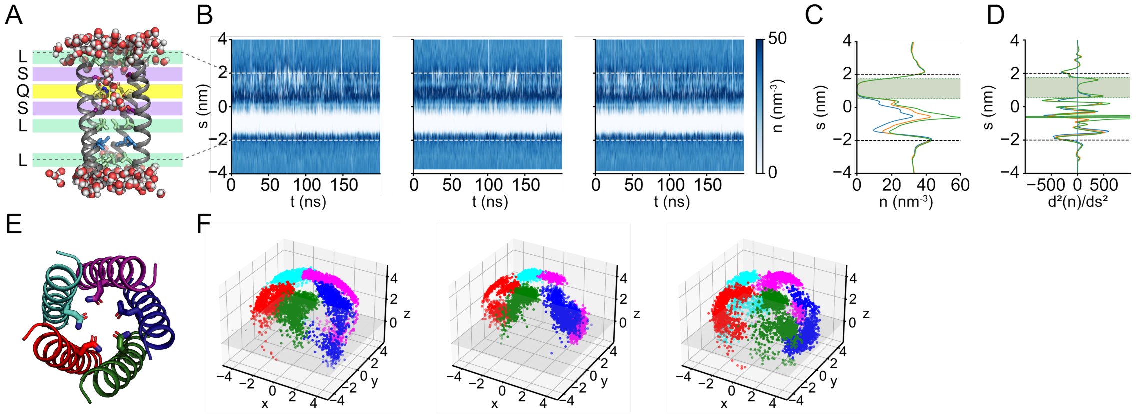

**Supplementary Figure 12. All MD simulation replicates for LQLL I6S/I13S.** (A) Representative snapshot of LQLL I6S/I13S. Waters within 3 Å of the protein are shown, and pore-facing residues are colored as in **Fig. 2**. (B) Water density profiles for all three production runs (1-3, left to right). Dashed lines represent the protein boundaries within the simulation. (C) Average water density line plots are shown for each production (1=blue, 2=orange, 3=green). (D) Second derivative of the average water density plot from (C). Shaded hydrophobic regions were determined by points where there is an intercept at x=0 within the protein boundaries (dashed). (E) Model pentamer with each of its five helices colored to denote the separate chains. (F) For all three production runs (1-3, left to right), $\vec{v}_{Gln}$ is plotted for the five Gln10 residues over the course of the simulation. The Cα plane is denoted in gray for (x, y, z = 0).

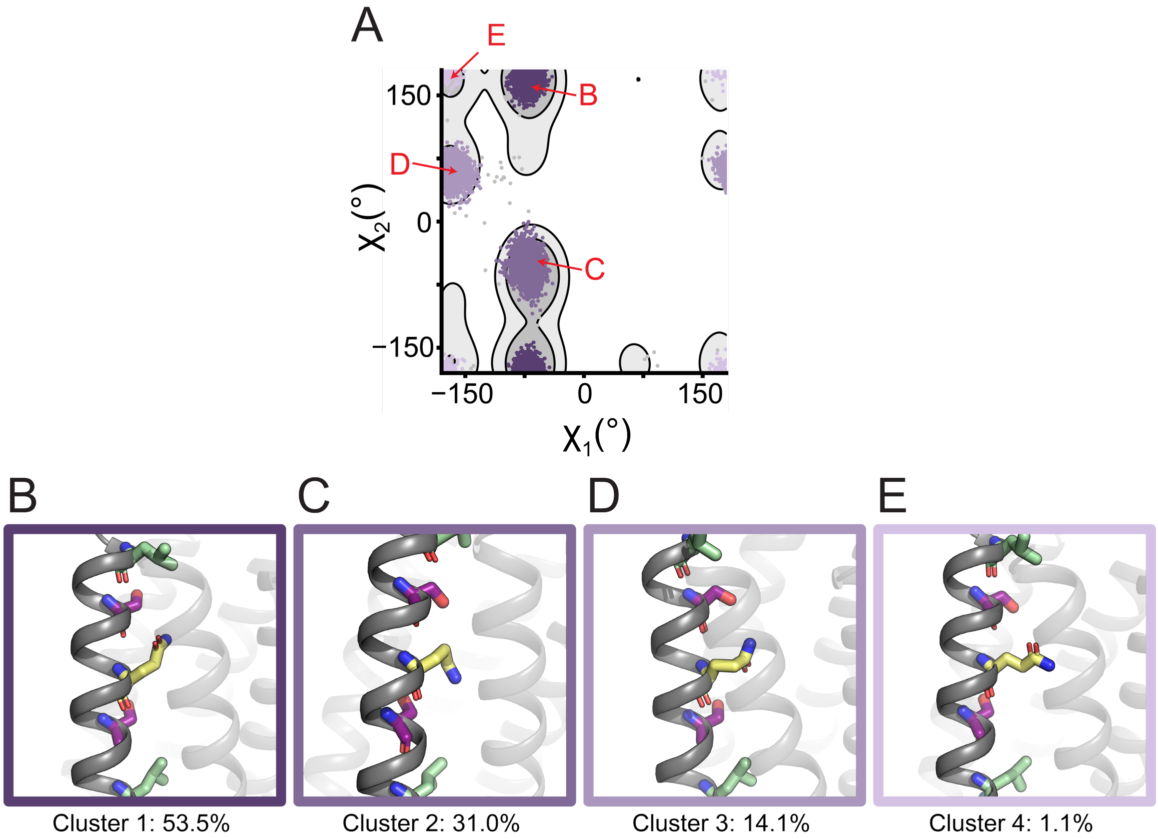

**Supplementary Figure 13. $\chi_1$/$\chi_2$ rotamer analysis of LQLL I6S/I13S MD simulations.** (A) Plot of $\chi_1$/$\chi_2$ dihedral angles from Gln sidechains sampled from MD simulations for each of the mutants. These are overlaid on a contour map of Gln rotameric states sampled from a curated structural database of transmembrane proteins, where lighter gray contours denote the top 70% and darker gray contours denote the top 90%. Simulation points are colored by cluster with more prevalent clusters represented in darker colors. (B-D) Representative snapshots from each cluster are shown with pore-facing residues colored as in **Fig. 2**. Boxes are colored by cluster and labeled on the graph (A).

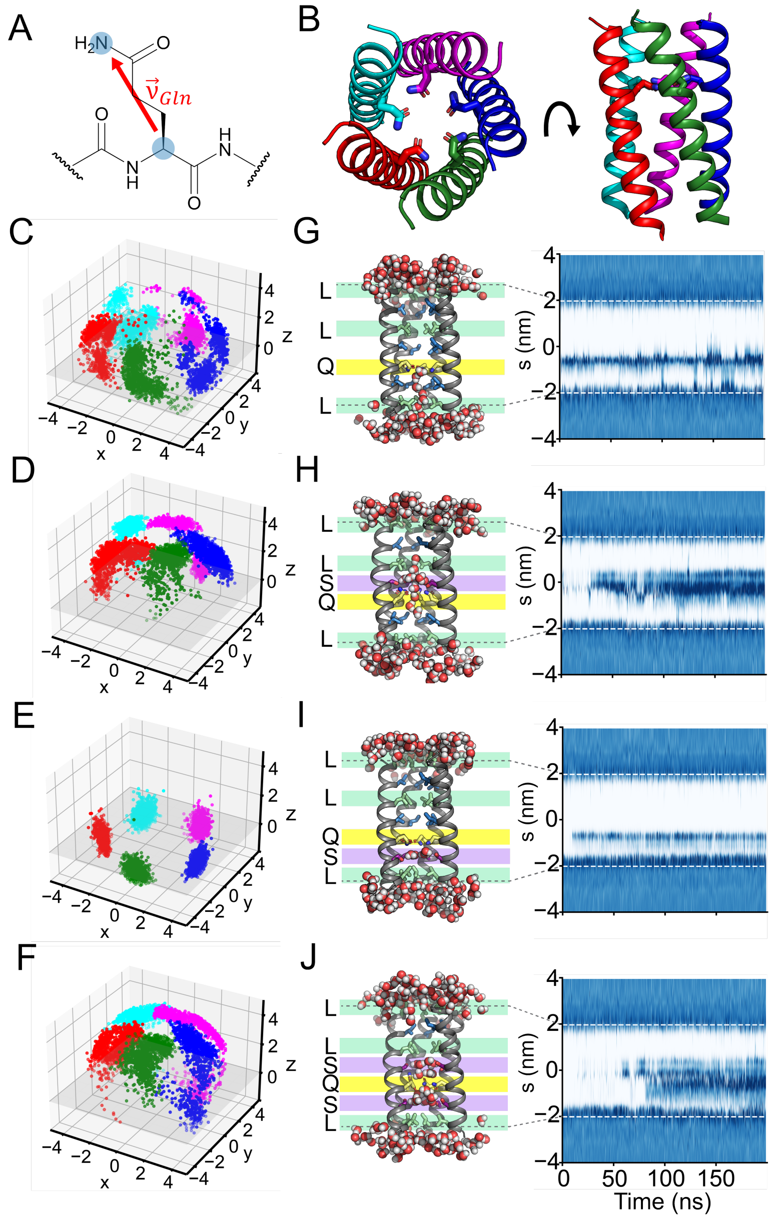

**Supplementary Figure 14. MD simulations indicate Ser substitutions affect Gln sidechain dynamics and pore hydration.** (A) $\vec{v}_{\mathrm{Gln}}$ is defined as the vector from the Cα to the Nε of the Gln sidechain. (B) Structure of LQLL (pdb 7UDZ) with each of its five helices colored to denote the separate chains. For each of the mutants (C) LLQL, (D) LLQL I13S, (E) LLQL I20S, and (F) LLQL I13S/I20S, $\vec{v}_{\mathrm{Gln}}$ is plotted for the five Gln17 residues over the course of the simulation. The Cα plane is denoted in gray for (x, y, z = 0). Representative snapshots and water density profiles from a single production run for (G) LLQL, (H) LLQL I13S, (I) LLQL I20S, and (J) LLQL I13S/I20S are shown. Waters within 3 Å of the protein are shown, and the pore-facing residues of interest are colored as in **Fig. 2**. Dashed lines represent the protein boundaries within the simulation.

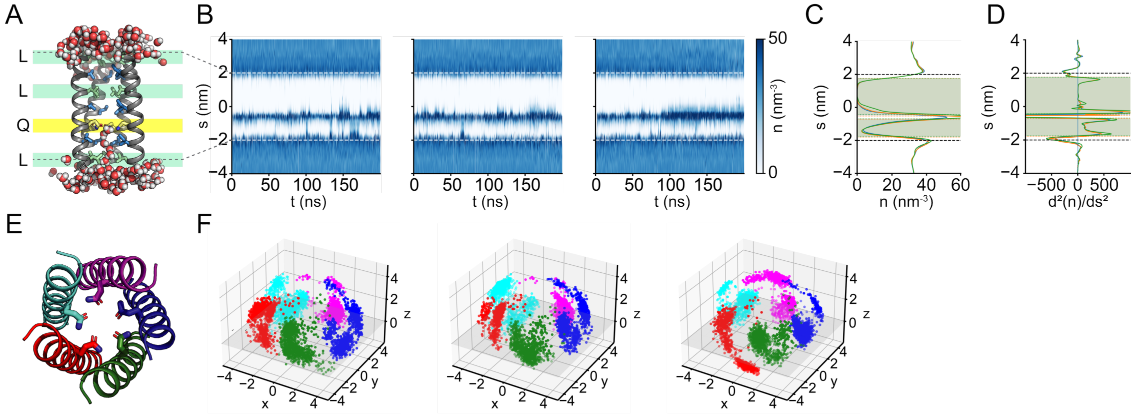

**Supplementary Figure 15. All MD simulation replicates for LLQL.** (A) Representative snapshot of LLQL. Waters within 3 Å of the protein are shown, and pore-facing residues are colored as in **Fig. 2**. (B) Water density profiles for all three production runs (1-3, left to right). Dashed lines represent the protein boundaries within the simulation. (C) Average water density line plots are shown for each production (1=blue, 2=orange, 3=green). (D) Second derivative of the average water density plot from (C). Shaded hydrophobic regions were determined by points where there is an intercept at x=0 within the protein boundaries (dashed). (E) Model pentamer with each of its five helices colored to denote the separate chains. (F) For all three production runs (1-3, left to right), $\vec{v}_{Gln}$ is plotted for the five Gln10 residues over the course of the simulation. The Cα plane is denoted in gray for (x, y, z = 0).

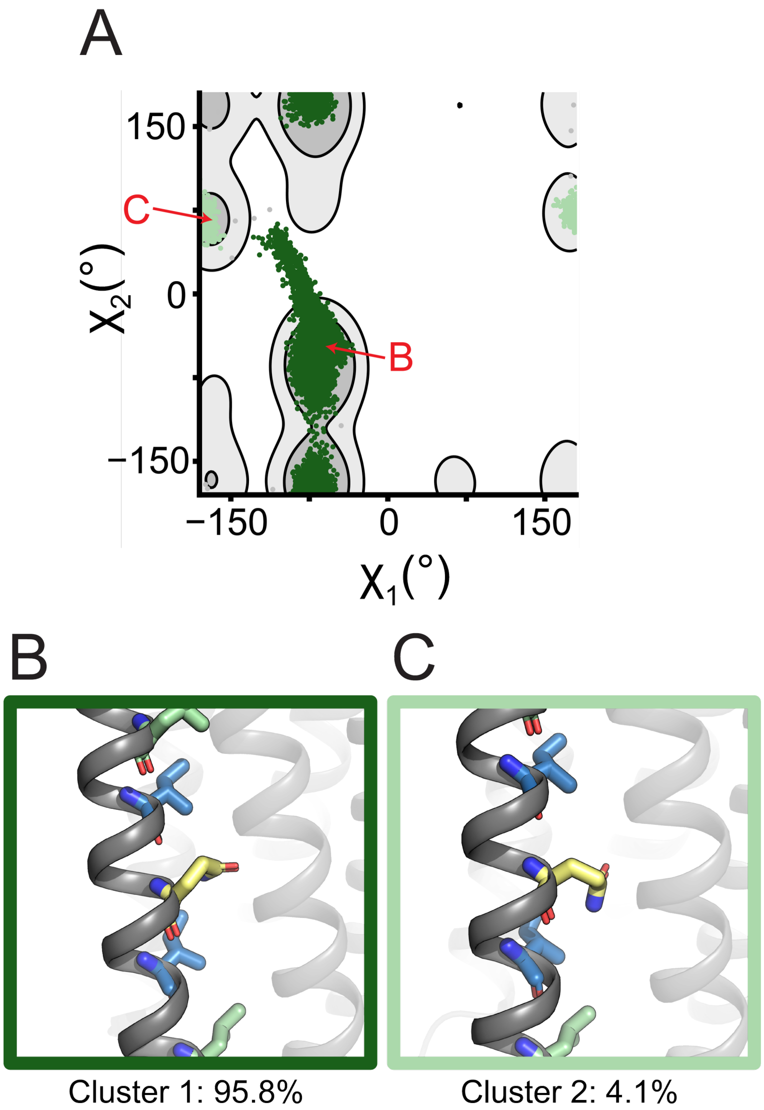

**Supplementary Figure 16. $\chi_1/\chi_2$ rotamer analysis of LLQL MD simulations.** (A) Plot of $\chi_1/\chi_2$ dihedral angles from Gln sidechains sampled from MD simulations for each of the mutants. These are overlaid on a contour map of Gln rotameric states sampled from a curated structural database of transmembrane proteins, where lighter gray contours denote the top 70% and darker gray contours denote the top 90%. Simulation points are colored by cluster with more prevalent clusters represented in darker colors. (B-D) Representative snapshots from each cluster are shown with pore-facing residues colored as in **Fig. 2**. Boxes are colored by cluster and labeled on the graph (A).

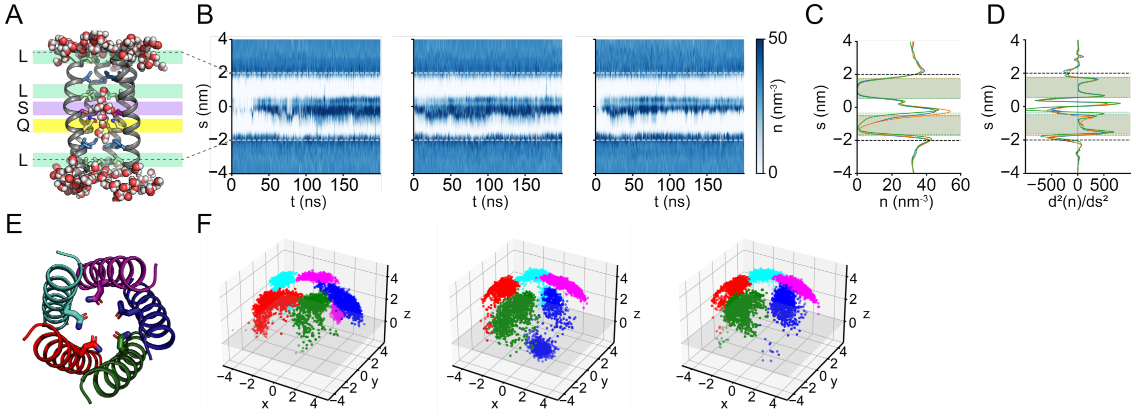

**Supplementary Figure 17. All MD simulation replicates for LLQL I13S.** (A) Representative snapshot of LLQL I13S. Waters within 3 Å of the protein are shown, and pore-facing residues are colored as in **Fig. 2**. (B) Water density profiles for all three production runs (1-3, left to right). Dashed lines represent the protein boundaries within the simulation. (C) Average water density line plots are shown for each production (1=blue, 2=orange, 3=green). (D) Second derivative of the average water density plot from (C). Shaded hydrophobic regions were determined by points where there is an intercept at x=0 within the protein boundaries (dashed). (E) Model pentamer with each of its five helices colored to denote the separate chains. (F) For all three production runs (1-3, left to right), $\vec{v}_{Gln}$ is plotted for the five Gln10 residues over the course of the simulation. The Cα plane is denoted in gray for (x, y, z = 0).

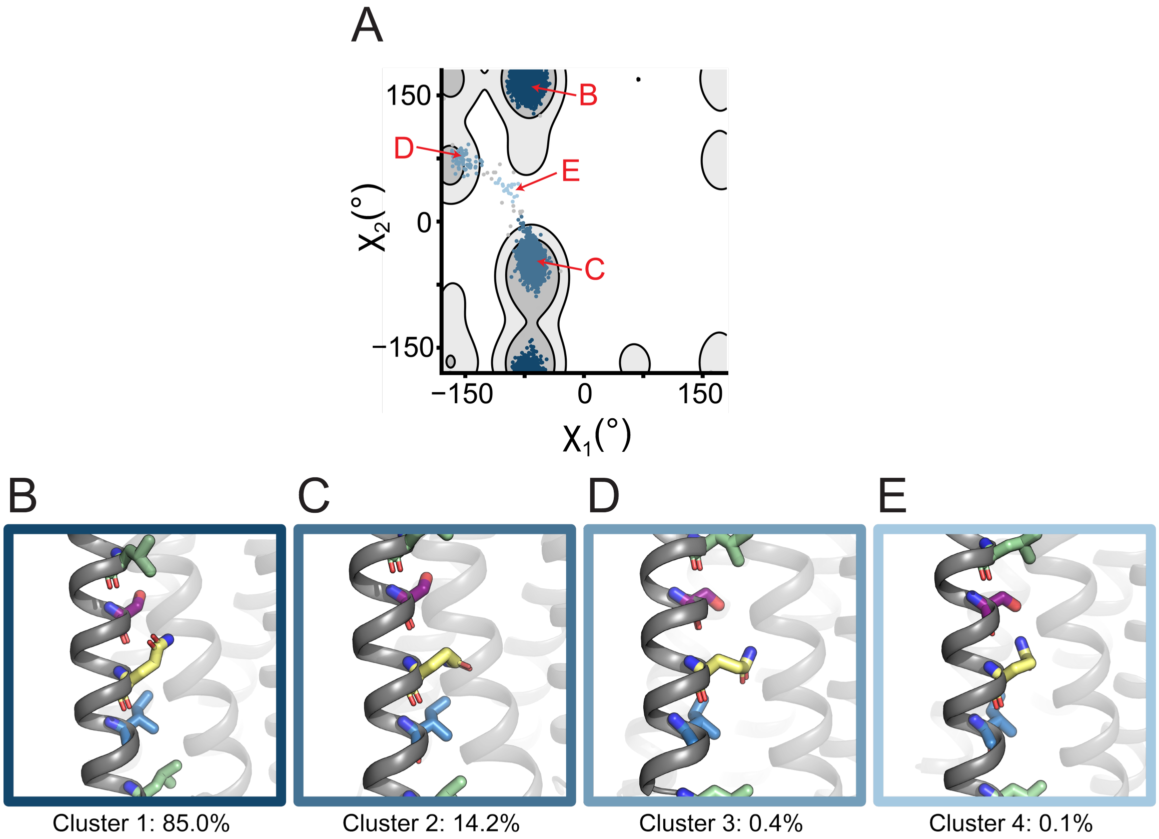

**Supplementary Figure 18. $\chi_1/\chi_2$ rotamer analysis of LLQL I13S MD simulations.** (A) Plot of $\chi_1/\chi_2$ dihedral angles from Gln sidechains sampled from MD simulations for each of the mutants. These are overlaid on a contour map of Gln rotameric states sampled from a curated structural database of transmembrane proteins, where lighter gray contours denote the top 70% and darker gray contours denote the top 90%. Simulation points are colored by cluster with more prevalent clusters represented in darker colors. (B-D) Representative snapshots from each cluster are shown with pore-facing residues colored as in **Fig. 2**. Boxes are colored by cluster and labeled on the graph (A).

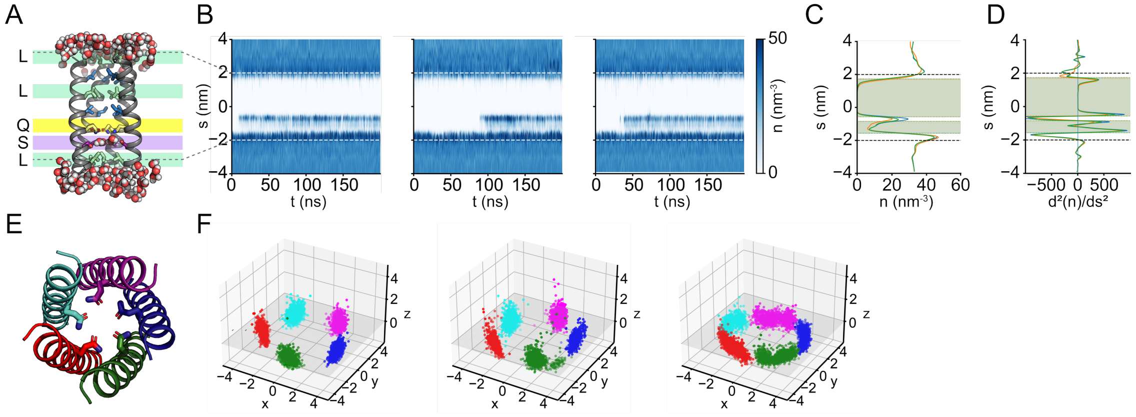

**Supplementary Figure 19. All MD simulation replicates for LLQL I20S.** (A) Representative snapshot of LLQL I20S. Waters within 3 Å of the protein are shown, and pore-facing residues are colored as in **Fig. 2**. (B) Water density profiles for all three production runs (1-3, left to right). Dashed lines represent the protein boundaries within the simulation. (C) Average water density line plots are shown for each production (1=blue, 2=orange, 3=green). (D) Second derivative of the average water density plot from (C). Shaded hydrophobic regions were determined by points where there is an intercept at x=0 within the protein boundaries (dashed). (E) Model pentamer with each of its five helices colored to denote the separate chains. (F) For all three production runs (1-3, left to right), $\vec{v}_{Gln}$ is plotted for the five Gln10 residues over the course of the simulation. The Cα plane is denoted in gray for (x, y, z = 0).

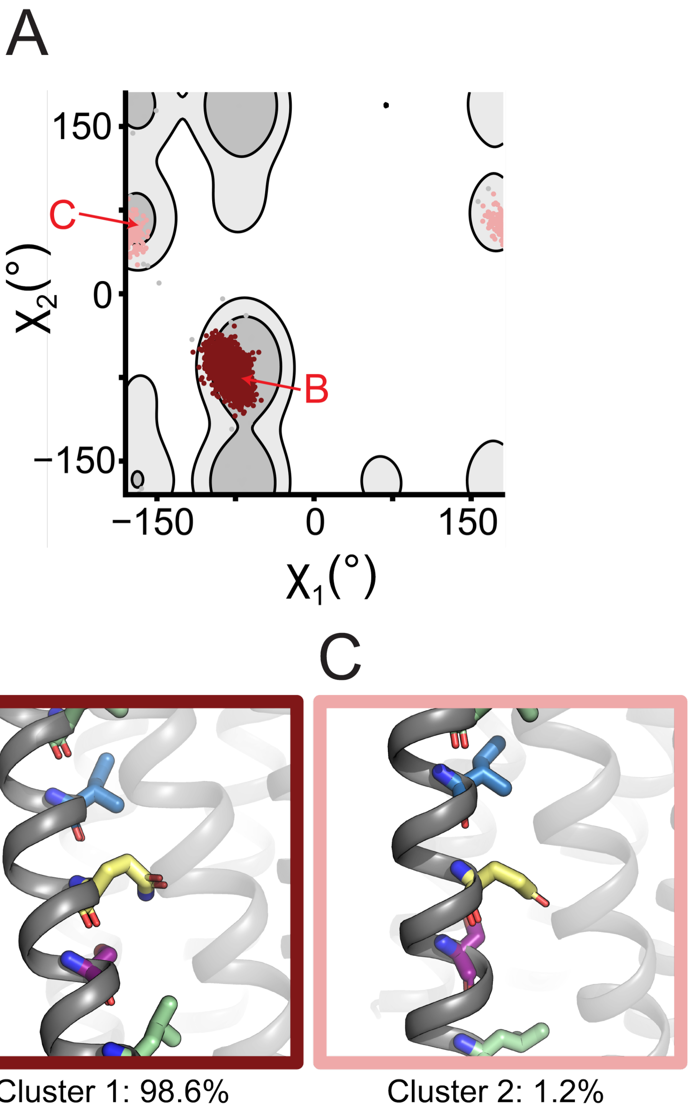

**Supplementary Figure 20. Supplementary Fig. S20. $\chi_1/\chi_2$ rotamer analysis of LLQL I20S MD simulations.** (A) Plot of $\chi_1/\chi_2$ dihedral angles from Gln sidechains sampled from MD simulations for each of the mutants. These are overlaid on a contour map of Gln rotameric states sampled from a curated structural database of transmembrane proteins, where lighter gray contours denote the top 70% and darker gray contours denote the top 90%. Simulation points are colored by cluster with more prevalent clusters represented in darker colors. (B-D) Representative snapshots from each cluster are shown with pore-facing residues colored as in **Fig. 2**. Boxes are colored by cluster and labeled on the graph (A).

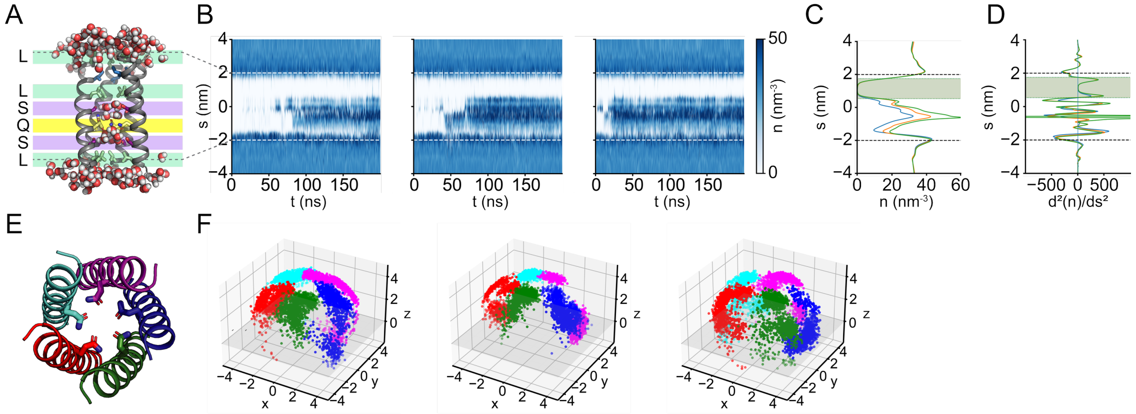

**Supplementary Figure 21. All MD simulation replicates for LLQL I13S/I20S.** (A) Representative snapshot of LLQL I13S/I20S. Waters within 3 Å of the protein are shown, and pore-facing residues are colored as in **Fig. 2**. (B) Water density profiles for all three production runs (1-3, left to right). Dashed lines represent the protein boundaries within the simulation. (C) Average water density line plots are shown for each production (1=blue, 2=orange, 3=green). (D) Second derivative of the average water density plot from (C). Shaded hydrophobic regions were determined by points where there is an intercept at x=0 within the protein boundaries (dashed). (E) Model pentamer with each of its five helices colored to denote the separate chains. (F) For all three production runs (1-3, left to right), $\vec{v}_{Gln}$ is plotted for the five Gln10 residues over the course of the simulation. The Cα plane is denoted in gray for (x, y, z = 0).

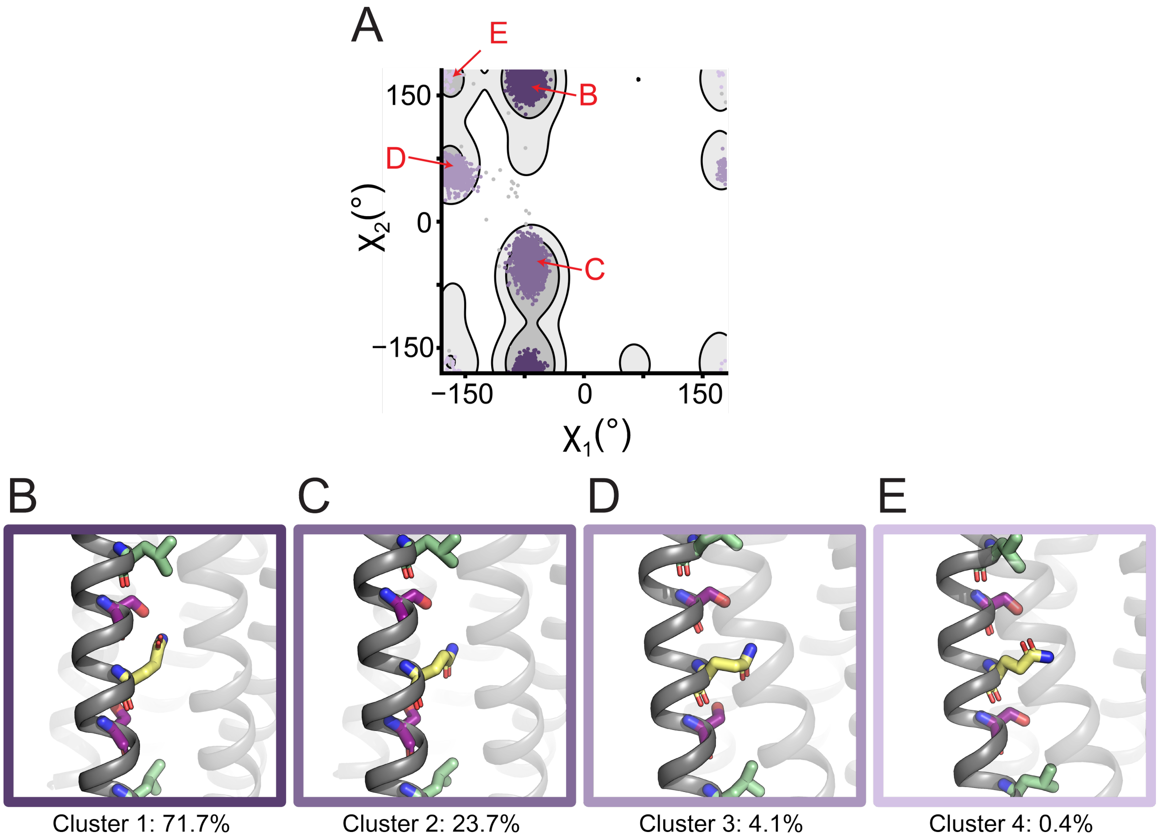

**Supplementary Figure 22. $\chi_1/\chi_2$ rotamer analysis of LLQL I13S/I20S MD simulations.** (A) Plot of $\chi_1/\chi_2$ dihedral angles from Gln sidechains sampled from MD simulations for each of the mutants. These are overlaid on a contour map of Gln rotameric states sampled from a curated structural database of transmembrane proteins, where lighter gray contours denote the top 70% and darker gray contours denote the top 90%. Simulation points are colored by cluster with more prevalent clusters represented in darker colors. (B-D) Representative snapshots from each cluster are shown with pore-facing residues colored as in **Fig. 2**. Boxes are colored by cluster and labeled on the graph (A).

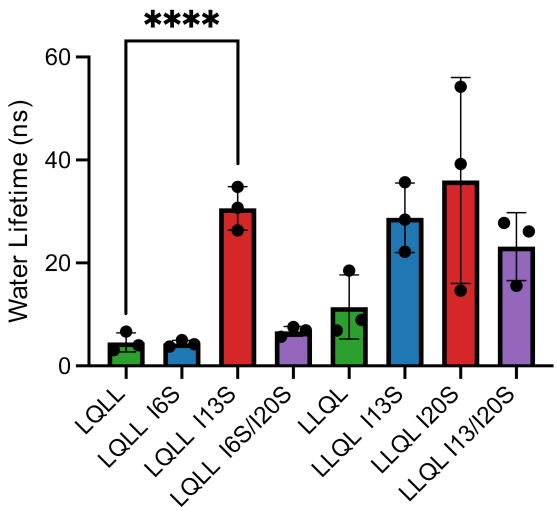

**Supplementary Figure 23. Water lifetimes for all variants across all 3 simulations.** Water lifetimes were calculated by tracking waters within a cylindrical region defining the pore. Waters were considered inside the pore if their coordinates fell within a cylinder defined by the Cα atoms of residues 5 and 22 across all five helices. For each water molecule entering this region, the time between entry and exit was measured as its pore residence time. Bars represent the mean value across n=3 simulations with error bars showing SEM. To compare significance amongst mutants within their families, a one-way ANOVA and post-hoc Dunnett's test was performed, and mutants were compared to the parent (LQLL or LLQL). Only LQLL I13S showed a significant difference from the parent with a p-value <0.0001 (**Supplementary Table 14**).

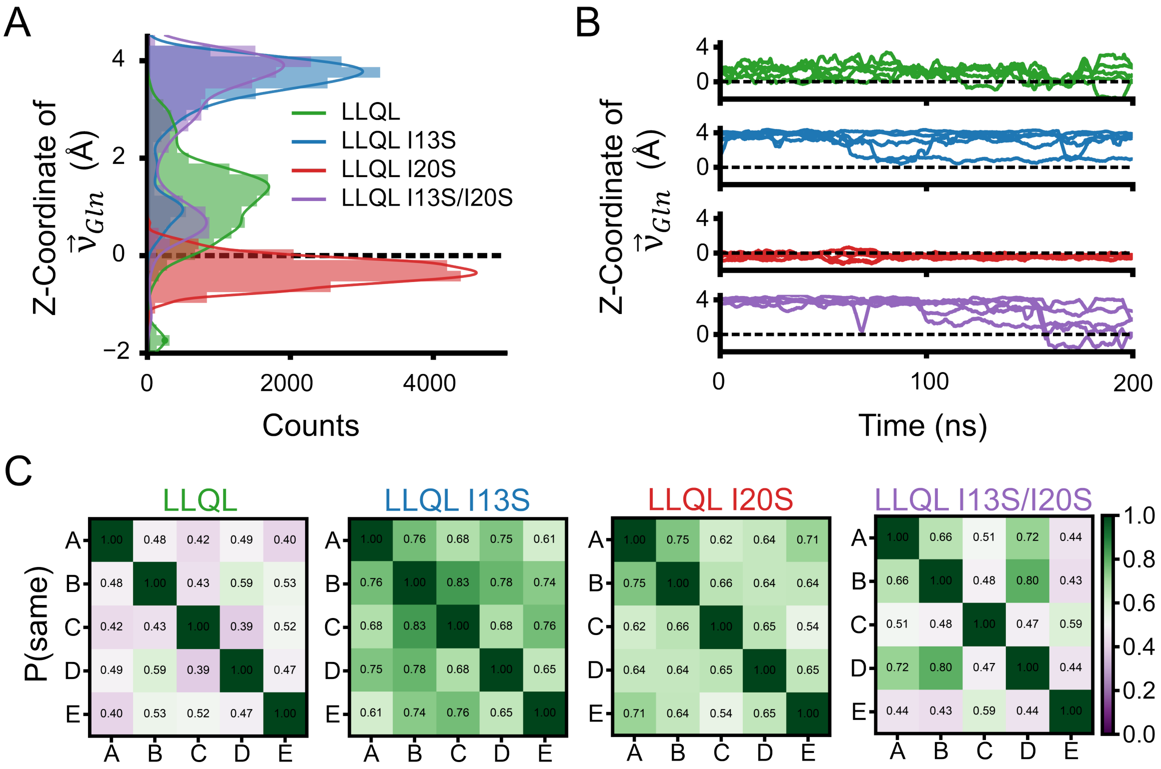

**Supplementary Figure 24. MD simulations of LLQL reveal double Ser mutant breaks channel symmetry.** (A) Histogram of z-coordinate of $\vec{v}_{Gln}$ for all Ser mutants of LLQL. Counts include all five Gln17 residues within the pore across the full simulation and all three replicates for a total of 15,000 data points per variant. The Cα plane is shown as a dashed line. (B) Line plots of the z-coordinate of each of the five pore-lining Gln17 residues as a function of time for a single simulation trajectory of each mutant. (C) Pairwise state agreement matrices for Gln sidechain in LLQL variants.

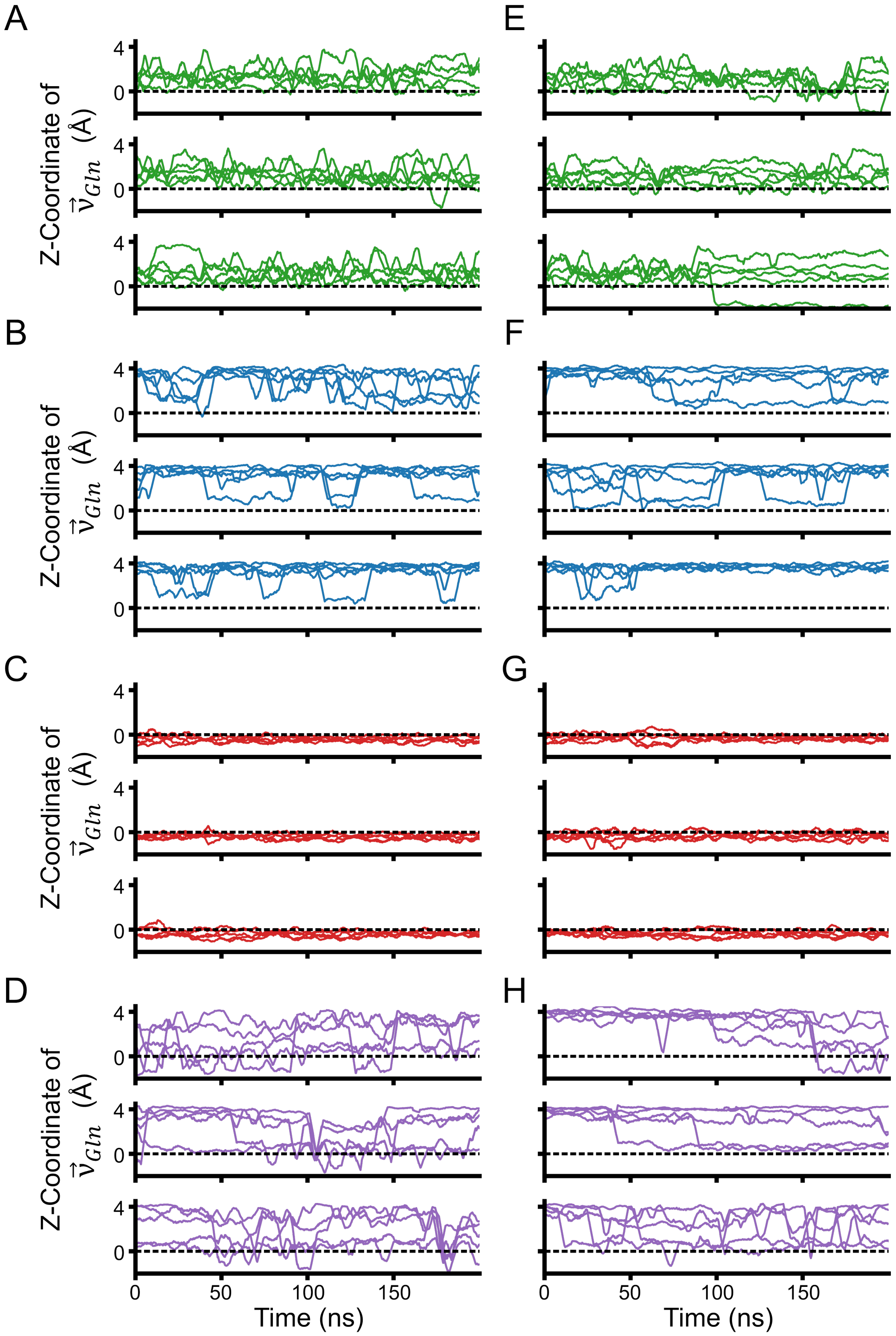

**Supplementary Figure 25. Gln vector z-coordinate for all simulations.** Line plots of the z-coordinate of each of the five pore-lining Gln 10 (A-D) and Gln17 (E-H) residues as a function of time for simulation trajectories 1-3 (top-bottom) of each mutant. Mutants shown are LQLL (A), LQLL I6S (B), LQLL I13S (C), LQLL I6S/I13S (D), LLQL (E), LLQL I13S (F), LLQL I20S (G), and LLQL I13S/I20S (H).

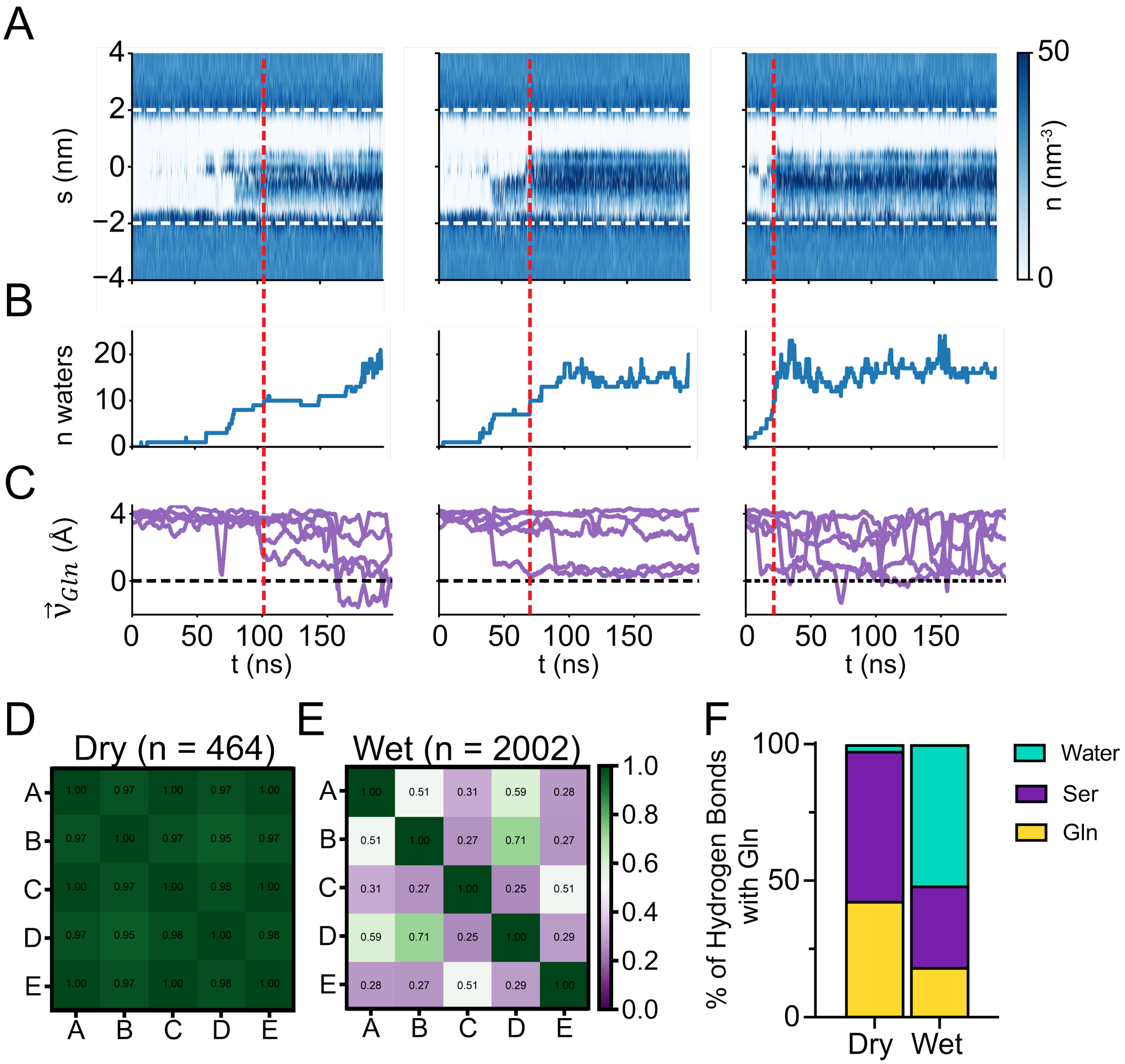

**Supplementary Figure 26. Comparison of hydrated and dehydrated frames of simulation for LLQL I13S/I20S.** (A-C) Replicate productions 1-3 (left to right) for LLQL I13S/I20S. Red dashed lines represent threshold cutoff of n=10 waters for a “hydrated” pore. (A) Water density profiles with dashed lines representing the protein boundaries within the simulation. (B) Number of waters within the pore during each frame of the simulation. (C) Line plots of the z-coordinate of each of the five pore-lining Gln17 residues as a function of time. The Cα plane is denoted by a dashed line. (D-E) Pairwise state agreement matrices for Gln sidechain for dry frames with n=0 or 1 water (D) and wet frames with all frames after the n=10 water threshold is met (E). (F) Percent hydrogen bonding partners for Gln17 in LLQL IdubS for the same dry vs wet frames from D and E.

**Supplementary Table 1. Peptide sequences and observed masses.** Peptides were synthesized via solid phase synthesis as described in materials and methods. Masses of synthesized peptides were observed using MALDI.

| Peptide | Sequence | Expected Mass (Da) | Observed Mass (Da) |
|---|---|---|---|
| AM2 TM | SSDPLVVAASIIGILHLILWILDRL | 2770.3 | 2768.5 |
| LLLL | DS $^{3}$LKWIVFL $^{10}$LFLIVLL $^{17}$LLAIVFL LRG | 3027.9 | 3027.9 |
| LQLL | DS $^{3}$LKWIVFL $^{10}$QFLIVLL $^{17}$LLAIVFL LRG | 3042.9 | 3041.8 |
| LQLL I6S | DS $^{3}$LKWSVFL $^{10}$QFLIVLL $^{17}$LLAIVFL LRG | 3016.8 | 3015.7 |
| LQLL I13S | DS $^{3}$LKWIVFL $^{10}$QFLSVLL $^{17}$LLAIVFL LRG | 3016.8 | 3015.7 |
| LQLL I20S | DS $^{3}$LKWIVFL $^{10}$QFLIVLL $^{17}$LLASVFL LRG | 3016.8 | 3015.9 |
| LQLL I6S/I13S | DS $^{3}$LKWSVFL $^{10}$QFLSVLL $^{17}$LLAIVFL LRG | 2990.7 | 2989.4 |
| LLQL | DS $^{3}$LKWIVFL $^{10}$LFLIVLL $^{17}$QLAIVFL LRG | 3042.9 | 3041.8 |
| LLQL I6S | DS $^{3}$LKWSVFL $^{10}$LFLIVLL $^{17}$QLAIVFL LRG | 3016.8 | 3015.7 |
| LLQL I13S | DS $^{3}$LKWIVFL $^{10}$LFLSVLL $^{17}$QLAIVFL LRG | 3016.8 | 3015.7 |
| LLQL I20S | DS $^{3}$LKWIVFL $^{10}$LFLIVLL $^{17}$QLASVFL LRG | 3016.8 | 3015.7 |
| LLQL I13S/I20S | DS $^{3}$LKWIVFL $^{10}$LFLSVLL $^{17}$QLASVFL LRG | 2990.7 | 2989.2 |

**Supplementary Table 2. Unpaired t-test between Empty and AM2 TM proton fluxes.** Proton fluxes between positive (AM2 TM) and negative (empty) controls (**Fig. 5C**) were compared using an unpaired t-test with Welch's correction to account for differing variances of data sets. Flux from liposomes containing the natural channel construct AM2-TM showed statistical significance compared to liposomes with no peptide incorporated.

| Empty vs AM2-TM: Unpaired t-test with Welch's correction | |
|---|---|
| **P value** | <0.0001 |
| **P value summary** | **** |
| **Significantly different (P < 0.05)?** | Yes |
| **One- or two-tailed P value?** | Two-tailed |
| **Welch-corrected t, df** | t=17.54, df=7.799 |

**Supplementary Table 3. Dunnett's test of all proton flux rates for mutants compared to LLLL.** Flux for all mutant peptides were compared to the fully hydrophobic construct LLLL(35) using an ordinary one-way ANOVA with post-hoc Dunnett's test. All mutant constructs showed significantly more proton flux compared to LLLL.

| Dunnett's Test | Mean diff. | 95.00% CI of diff. | Below Threshold? | Summary | Adjusted p-value |
|---|---|---|---|---|---|
| **LLLL vs LQLL** | -7.963e-006 | -1.521e-005 to -7.163e-007 | Yes | * | 0.0249 |
| **LLLL vs LQLL I6S** | -1.184e-005 | -1.909e-005 to -4.593e-006 | Yes | *** | 0.0003 |
| **LLLL vs LQLL I13S** | -1.228e-005 | -1.953e-005 to -5.037e-006 | Yes | *** | 0.0001 |
| **LLLL vs LQLL I6S/I13S** | -2.769e-005 | -3.493e-005 to -2.044e-005 | Yes | **** | <0.0001 |
| **LLLL vs LLQL** | -7.666e-006 | -1.491e-005 to -4.190e-007 | Yes | * | 0.0335 |
| **LLLL vs LLQL I13S** | -1.323e-005 | -2.047e-005 to -5.980e-006 | Yes | **** | <0.0001 |
| **LLLL vs LLQL I20S** | -8.567e-006 | -1.581e-005 to -1.321e-006 | Yes | * | 0.0133 |
| **LLLL vs LLQL I13S/I20S** | -3.663e-005 | -4.387e-005 to -2.938e-005 | Yes | **** | <0.0001 |

**Supplementary Table 4. Tukey's multiple comparison test for all proton flux rates in the LQLL family.** Within the LQLL family of mutants (**Fig. 5D**), proton fluxes were compared using an ordinary one-way ANOVA with post-hoc Tukey's test to make multiple comparisons between mutants. Only the double serine mutant LQLL I6S/I13S showed statistical significance compared to all other mutants.

| Tukey's Test LQLL | Mean diff. | 95.00% CI of diff. | Below Threshold? | Summary | Adjusted p-value |
|---|---|---|---|---|---|
| **LQLL vs LQLL I6S** | -3.876e-006 | -1.050e-005 to 2.743e-006 | No | ns | 0.3955 |
| **LQLL vs LQLL I13S** | -4.321e-006 | -1.094e-005 to 2.298e-006 | No | ns | 0.3027 |
| **LQLL vs LQLL I6S/I13S** | -1.972e-005 | -2.634e-005 to -1.310e-005 | Yes | **** | <0.0001 |
| **LQLL I6S vs LQLL I13S** | -4.447e-007 | -7.064e-006 to 6.175e-006 | No | ns | 0.9978 |
| **LQLL I6S vs LQLL I6S/I13S** | -1.585e-005 | -2.247e-005 to -9.228e-006 | Yes | **** | <0.0001 |
| **LQLL I13S vs LQLL I6S/I13S** | -1.540e-005 | -2.202e-005 to -8.784e-006 | Yes | **** | <0.0001 |

**Supplementary Table 5. Tukey's multiple comparison test for all proton flux rates in the LLQL family.** Within the LLQL family of mutants (**Fig. 5E**), proton fluxes were compared using an ordinary one-way ANOVA with post-hoc Tukey's test to make multiple comparisons between mutants. Only the double serine mutant LLQL I13S/I20S showed statistical significance compared to all other mutants.

| Tukey's Test LLQL | Mean diff. | 95.00% CI of diff. | Below Threshold? | Summary | Adjusted p-value |
|---|---|---|---|---|---|
| **LLQL vs LLQL I13S** | -5.561e-006 | -1.415e-005 to 3.024e-006 | No | ns | 0.3092 |
| **LLQL vs LLQL I20S** | -9.015e-007 | -9.486e-006 to 7.683e-006 | No | ns | 0.9916 |
| **LLQL vs LLQL I13S/I20S** | -2.896e-005 | -3.755e-005 to -2.038e-005 | Yes | **** | <0.0001 |
| **LLQL I13S vs LLQL I20S** | 4.659e-006 | -3.926e-006 to 1.324e-005 | No | ns | 0.4615 |
| **LLQL I13S vs LLQL I13S/I20S** | -2.340e-005 | -3.199e-005 to -1.482e-005 | Yes | **** | <0.0001 |
| **LLQL I20S vs LLQL I13S/I20S** | -2.806e-005 | -3.665e-005 to -1.948e-005 | Yes | **** | <0.0001 |

**Supplementary Table 6. Pore hydrophobic lengths.** Hydrophobic lengths obtained from MD simulations for all mutants.

| Design | Hydrophobic Length 1 (Å) | Hydrophobic Length 2 (Å) |
|---|---|---|
| LQLL | 20.8 ± 0.1 | 11.4 ± 0.5 |
| LQLL I6S | 22.9 ± 0.4 | |
| LQLL I13S | 14.1 ± 0.6 | 12.5 ± 0.2 |
| LQLL I6S/I13S | 14.6 ± 0.0 | |
| LLQL | 22.3 ± 0.4 | 10.6 ± 0.2 |
| LLQL I13S | 12.5 ± 0.3 | 11.9 ± 0.3 |
| LLQL I20S | 22.9 ± 0.1 | 7.3 ± 0.1 |
| LLQL I13S/I20S | 12.1 ± 0.1 | |

**Supplementary Table 7. Percent hydrogen bonding partners for Gln residues.** For each mutant all hydrogen bonds to Gln10 (LQLL family) or Gln17 (LLQL family) were extracted and pooled from all three production runs. Percentages are also represented graphically in **Supplementary Figure S5**.

| | Gln | Ser | Water |
|---|---|---|---|
| **LQLL** | 69.6 | | 30.4 |
| **LQLL I6S** | 27.6 | 42.2 | 30.2 |
| **LQLL I13S** | 83.4 | 0.1 | 16.5 |
| **LQLL I6S/I13S** | 13.2 | 29.5 | 57.3 |
| **LLQL** | 64.2 | | 35.8 |
| **LLQL I13S** | 33.0 | 38.2 | 28.8 |
| **LLQL I20S** | 86.8 | 0.0 | 13.2 |
| **LLQL I13S/I20S** | 24.0 | 35.8 | 40.2 |

**Supplementary Table 8. Gln $\chi_1/\chi_2$ cluster percentages and average ΔZ for each cluster.** The percent of frames for each cluster present in **Supplementary Figures S7, S9, S11, S13, S16, S18, S22, and S22** are shown. The average ΔZ values for the Gln in each cluster were also determined.

| | Cluster 1 | | Cluster 2 | | Cluster 3 | | Cluster 4 | |
|---|---|---|---|---|---|---|---|---|
| | % | ΔZ | % | ΔZ | % | ΔZ | % | ΔZ |
| **LQLL** | 75.2 | 0.77 | 24.2 | 2.73 | 0.4 | -1.25 | | |
| **LQLL I6S** | 81.1 | 3.55 | 17.2 | 1.09 | 1.4 | 1.85 | 0.2 | 1.60 |
| **LQLL I13S** | 99.0 | -0.39 | 0.9 | -0.80 | | | | |
| **LQLL I6S/I13S** | 53.5 | 3.21 | 31.0 | 0.60 | 14.1 | -0.33 | 1.1 | 1.76 |
| **LLQL** | 95.8 | 1.19 | 4.1 | -1.73 | | | | |
| **LLQL I13S** | 85.0 | 3.61 | 14.2 | 0.90 | 0.4 | 1.95 | 0.1 | 2.44 |
| **LLQL I20S** | 98.6 | -0.33 | 1.2 | -0.89 | | | | |
| **LLQL I13S/I20S** | 71.7 | 3.54 | 23.7 | 0.73 | 4.1 | -0.23 | 0.4 | 1.71 |

**Supplementary Table 9. Gln $\chi_1$/$\chi_2$ dihedrals for all crystal structures in Fig. 5.** For each crystal structure, the $\chi_1$/$\chi_2$ dihedrals were extracted for all Gln residues present. The models for LQLL, LLQL I13S, and LQLL I6S/I13S all have two pentameric bundles in the ASU resulting in 10 different Gln residues. The model for LQLL I13S only has one pentameric bundle in the ASU resulting in 5 different Gln residues.

| | Chain A | | Chain B | | Chain C | | Chain D | | Chain E | |
|---|---|---|---|---|---|---|---|---|---|---|
| | $\chi_1$ | $\chi_2$ | $\chi_1$ | $\chi_2$ | $\chi_1$ | $\chi_2$ | $\chi_1$ | $\chi_2$ | $\chi_1$ | $\chi_2$ |
| **LQLL** | -75.0 | -81.5 | -79.8 | -74.0 | -72.5 | -154.6 | -77.6 | -80.0 | -70.5 | -85.2 |
| | -77.7 | -74.1 | -75.3 | -88.0 | -65.0 | -107.1 | -79.8 | -76.9 | -69.5 | -85.9 |
| **LLQL I13S** | -80.8 | 152.0 | -80.2 | 152.6 | -80.1 | 152.3 | -81.3 | 151.6 | -79.8 | 154.3 |
| | -80.9 | 152.1 | -80.4 | 152.0 | -80.5 | 152.1 | -81.1 | 154.2 | -79.7 | 152.4 |
| **LQLL I13S** | -92.2 | -52.9 | -148.5 | 50.8 | -92.1 | -55.8 | -91.6 | -58.8 | -91.7 | -57.9 |
| **LQLL I6S/I13S** | -83.0 | -54.5 | -78.2 | 173.4 | -81.5 | 174.4 | -82.3 | -46.0 | -79.1 | 173.6 |
| | -78.8 | 172.5 | -80.3 | 173.5 | -82.5 | 175.3 | -80.3 | 174.7 | -145.7 | 42.7 |

**Supplementary Table 10. Channel crystallization conditions.** Solutions used for protein crystallography using lipidic cubic phase methods as described in Materials and Methods. Cryoprotectant was used as noted during the looping stage.

| Design | Condition | Cryoprotectant |
|---|---|---|
| **LQLL I13S** | 0.125 M CaCl2, 0.02 M TRIS pH 7.4, 30% PEG 3K | 30% PEG 400 |
| **LLQL I13S** | 6% EG, 0.1 M NaCacod pH 6, 6.6% PEG 8K, 0.15 M ZnAcet | N/A |
| **LQLL IdubS** | 0.1M NaCl, 0.1M HEPES pH 7.5, 12% PEG 4K | 30% PEG 400 |

**Supplementary Table 11. Crystallography data collection and refinement statistics for all structures.** All structures were obtained from single protein crystals. Statistics for reflections in the outermost shell are included in parentheses. All data represent isotropic statistics.

| Property | LQLL I13S | LLQL I13S | LQLL IdubS |
|---|---|---|---|
| **Data Collection** | | | |
| Space Group | P 42 21 2 | P 21 21 21 | P1 |
| **Cell Dimensions** | | | |
| a, b, c (Å) | 54.65, 54.65, 80.67 | 41.18, 85.41, 87.19 | 40.86, 45.09, 47.51 |
| a, b, g (°) | 90.00, 90.00, 90.00 | 90.00, 90.00, 90.00 | 117.73, 99.48, 106.02 |
| Resolution (Å) | 45.24-2.90 (3.08-2.90) | 61.01-2.24 (2.31-2.24) | 39.40-1.87 (1.91-1.87) |
| $R_{merge}$ | 0.211 (1.209) | 0.17 (2.53) | 0.11 (5.67) |
| <I/sI> | 6.8 (2.1) | 4.8 (0.5) | 5.6 (0.2) |
| Completeness (%) | 93.2 (100) | 92.7 (81.3) | 81.9 (64.9) |
| CC1/2 | 0.999 (0.836) | 1.00 (0.41) | 1.00 (0.12) |
| Redundancy | 11.8 (11.2) | 6.5 (6.5) | 3.8 (3.4) |
| **Refinement** | | | |
| Resolution (Å) | 2.90 | 2.80 | 2.25 |
| No. Reflections | 2796 | 7379 | 11009 |
| $R_{work}$/ $R_{free}$ | 0.244, 0.278 | 0.242, 0.280 | 0.207, 0.247 |
| **No. Atoms** | | | |
| Protein | 1012 | 2116 | 2067 |
| Ligand | 14 | 48 | 23 |
| Water | 5 | 9 | 1 |
| **B-Factors** | | | |
| Protein | 52.9 | 48.0 | 38.3 |
| Ligand/Ion | 54.1 | 76.5 | 45.7 |
| Water | 30.5 | 57.8 | 47.4 |
| **R.M.S. deviations** | | | |
| Bond Lengths (Å) | 0.0061 | 0.0080 | 0.0057 |
| Bond Angles (°) | 1.5850 | 1.8290 | 1.2400 |
| **Ramachandran Statistics** | | | |
| Outliers (%) | 0 | 0 | 0 |
| Allowed (%) | 0.9 | 1.7 | 1.3 |
| Favored (%) | 99.1 | 98.3 | 98.7 |

**Supplementary Table 12. Tukey's multiple comparison test for all hydrophobic lengths in the LQLL family.** Within the LQLL family of mutants (**Supplementary Figure S3**), proton fluxes were compared using an ordinary one-way ANOVA with post-hoc Tukey's test to make multiple comparisons between mutants. The mutants with decreased hydrophobic lengths (LQLL I13S and LQLL I6S/I13S) showed no significant difference.

| Tukey's Test LQLL | Mean diff. | 95.00% CI of diff. | Below Threshold? | Summary | Adjusted p-value |
|---|---|---|---|---|---|
| **LQLL vs LQLL I6S** | -2.035 | -3.568 to -0.5017 | Yes | * | 0.0119 |
| **LQLL vs LQLL I13S** | 6.722 | 5.189 to 8.255 | Yes | **** | <0.0001 |
| **LQLL vs LQLL I6S/I13S** | 6.195 | 4.662 to 7.728 | Yes | **** | <0.0001 |
| **LQLL I6S vs LQLL I13S** | 8.756 | 7.223 to 10.29 | Yes | **** | <0.0001 |
| **LQLL I6S vs LQLL I6S/I13S** | 8.230 | 6.697 to 9.763 | Yes | **** | <0.0001 |
| **LQLL I13S vs LQLL I6S/I13S** | -0.5264 | -2.059 to 1.007 | No | ns | 0.6997 |

**Supplementary Table 13. Tukey's multiple comparison test for all hydrophobic lengths in the LLQL family.** Within the LLQL family of mutants (**Supplementary Figure S3**), proton fluxes were compared using an ordinary one-way ANOVA with post-hoc Tukey's test to make multiple comparisons between mutants. The mutants with decreased hydrophobic lengths (LLQL I13S and LLQL I13S/I20S) showed no significant difference.

| Tukey's Test LLQL | Mean diff. | 95.00% CI of diff. | Below Threshold? | Summary | Adjusted p-value |
|---|---|---|---|---|---|
| **LLQL vs LLQL I13S** | 9.780 | 8.714 to 10.85 | Yes | **** | <0.0001 |
| **LLQL vs LLQL I20S** | -0.5919 | -1.658 to 0.4737 | No | ns | 0.3486 |
| **LLQL vs LLQL I13S/I20S** | 10.16 | 9.093 to 11.22 | Yes | **** | <0.0001 |
| **LLQL I13S vs LLQL I20S** | -10.37 | -11.44 to -9.306 | Yes | **** | <0.0001 |
| **LLQL I13S vs LLQL I13S/I20S** | 0.3787 | -0.6869 to 1.444 | No | ns | 0.6783 |
| **LLQL I20S vs LLQL I13S/I20S** | 10.75 | 9.684 to 11.82 | Yes | **** | <0.0001 |

**Supplementary Table 14. Dunnett's test of all water lifetimes for mutants compared to their parents.** Water lifetimes for all mutant peptides were compared to their respective parent scaffolds (LQLL and LLQL) using an ordinary one-way ANOVA with post-hoc Dunnett's test. Only LQLL I13S showed a significant difference compared to LQLL.

| Dunnett's Test | Mean diff. | 95.00% CI of diff. | Below Threshold? | Summary | Adjusted p-value |
|---|---|---|---|---|---|
| **LQLL vs LQLL I6S** | 0.2323 | -5.361 to 5.826 | No | ns | 0.9987 |
| **LQLL vs LQLL I13S** | -26.01 | -31.61 to -20.42 | Yes | **** | <0.0001 |
| **LQLL vs LQLL I6S/I13S** | -2.142 | -7.735 to 3.451 | No | ns | 0.5800 |
| **LLQL vs LLQL I13S** | -17.33 | -44.36 to 9.702 | No | ns | 0.2275 |
| **LLQL vs LLQL I20S** | -24.59 | -51.62 to 2.442 | No | ns | 0.0736 |
| **LLQL vs LLQL I13S/I20S** | -11.74 | -38.77 to 15.29 | No | ns | 0.4925 |
